# A Single-Cell Temporal Atlas of Mouse Nasal Embryonic Development

**DOI:** 10.64898/2026.02.15.705966

**Authors:** Huan Chen, Yingxiu Chen, Mengjie Pan, Ziyu Feng, Baomei Cai, Yiyi Cheng, Sihao Chen, Jiehong Deng, Xia Yao, Chunhua Zhou, Yunjing Du, Wei He, Ruifang Zhang, Yudong Fu, Shujuan Liu, Lihui Lin, Shengyong Yu, Yuehong Yan, Duanqing Pei, Dajiang Qin, Jiekai Chen, Shangtao Cao

## Abstract

The mammals breathe and perceive odors through anatomically and functionally distinct respiratory and olfactory regions of the nose; however, how the early embryo endows this system remains largely unknown. Here, we present a temporal high-resolution single-cell RNA sequencing (scRNA-seq) atlas of mouse nasal development, comprising 183,247 cells collected from 20 embryos spanning embryonic day 10.5 (E10.5) to E18.5. We identified 7 major cell clusters and 52 molecularly defined subtypes, including previously unrecognized rare populations and niche-derived signaling components involved in progenitor cell maintenance. Lineage reconstruction of epithelial and mesenchymal compartments revealed dynamic cell fate transitions and highlighted candidate regulators of lineage specification, including *Foxa1* as a regulator of nasal respiratory ciliated epithelium differentiation. Integration with adult nasal epithelium datasets further delineated transcriptional dynamics of olfactory stem cells. This study provides a molecular roadmap of nasal development and offers a valuable resource for future research into nasal biology and regenerative strategies.

## Introduction

The nose is a principal respiratory and sensory organ in vertebrates, primarily responsible for air filtration, olfaction, and immune defense. Over the past decades, significant progress has been made in understanding nasal development from anatomical and histological perspectives (Cuschieri and Bannister, 1975; Kim et al., 2004; Sokpor et al., 2018; Kim et al., 2023; Parslow et al., 2024). More recently, single-cell technologies have enabled the construction of cellular atlases of the adult nasal epithelium or individual developmental stage, with most studies focusing on the olfactory region (Tsukahara et al., 2021; Horgue et al., 2022; Amato et al., 2024; Hills et al., 2024; Ualiyeva et al., 2024; Yang et al., 2024; Taroc et al., 2025). However, the molecular and cellular mechanisms underlying embryonic nasal development remain largely unclear, particularly how diverse cell populations coordinate to shape nasal structures during early development.

Embryonic nasal development is a complex and tightly regulated process. In mice, as in most mammals, the nasal placode arises from the frontonasal process and undergoes sequential morphogenesis through intricate epithelial–mesenchymal interactions (LaMantia et al., 2000; Streit, 2004; Balmer and LaMantia, 2005; Chen et al., 2009; Maier et al., 2011; Forni et al., 2013; Cho et al., 2019). The surface ectoderm thickens and invaginates into the underlying mesenchyme, forming the nasal pit by embryonic day 10.5 (E10.5), which gives rise to the primitive respiratory and olfactory epithelium and forms cavities through repeated tissue folding. Concurrently, mesoderm and neural crest–derived mesenchymal cells differentiate to shape the nasal framework via chondrogenesis and osteogenesis (Katoh et al., 2011; Suzuki and Osumi, 2015). These events are orchestrated by multiple conserved signaling pathways, including FGF, BMP, WNT, SHH and TGF-β, which regulate regional patterning and cell fate specification (Gong et al., 2009; Gokoffski, 2010; Maier et al., 2010; Maier et al., 2011; Forni et al., 2013; Zhu et al., 2016). Despite these well-characterized morphogenetic events, the detailed cellular composition, dynamic changes, and molecular regulation during the embryonic stages remain incompletely understood. In particular, the identification and characterization of nasal stem and progenitor cells, as well as the regulatory mechanisms guiding their lineage commitment, especially toward the respiratory epithelium, are still poorly defined.

To address these gaps, we generated a high-resolution single-cell RNA sequencing (scRNA-seq) temporal atlas of mouse nasal development during embryogenesis. This dataset allowed us to uncover previously unrecognized cell subtypes, novel marker genes, and niche-derived factors involved in progenitor cell maintenance. By reconstructing lineage trajectories of both epithelial and mesenchymal compartments, we identified key regulators governing cell fate decisions, notably highlighting the role of *Foxa1* in specifying the ciliated lineage of nasal respiratory epithelium. Furthermore, by integrating our dataset with published adult nasal epithelium atlases, we systematically traced the transcriptional dynamics of globose basal cells within the olfactory epithelium. Together, our study provides a molecular and cellular roadmap of mouse nasal development at single-cell resolution. It not only advances fundamental knowledge of nasal morphogenesis but also establishes a valuable resource for future studies on gene regulation, congenital nasal disorders, and regenerative medicine.

## Results

### scRNA-seq atlas of mouse nasal embryonic development

To systematically characterize the cellular composition and transcriptional dynamics underlying nasal morphogenesis at single-cell resolution, we collected mouse embryos from embryonic day 10.5 (E10.5) to E18.5 at 24-h intervals, and performed precise microdissection of nasal tissues, with at least two biological replicates for each stage (Figs. 1A and S1A). Single-cell suspensions were processed using the 10x Genomics platform, and after quality control, we obtained a developmental dataset comprising 183,247 high-quality cells, with a median of 3,777 detected genes and 12,822 UMIs per cell (Fig. S1B and S1C; Table S1). Uniform Manifold Approximation and Projection (UMAP) revealed a continuous trajectory of cellular states that recapitulated developmental progression (Fig. S1D). Pseudo–bulk principal component analysis (PCA) further demonstrated a robust temporal ordering along the first principal component (PC1), closely aligned with the actual developmental timeline, supporting the robustness and reproducibility of the dataset (Fig. S1E).

**Fig. 1.**
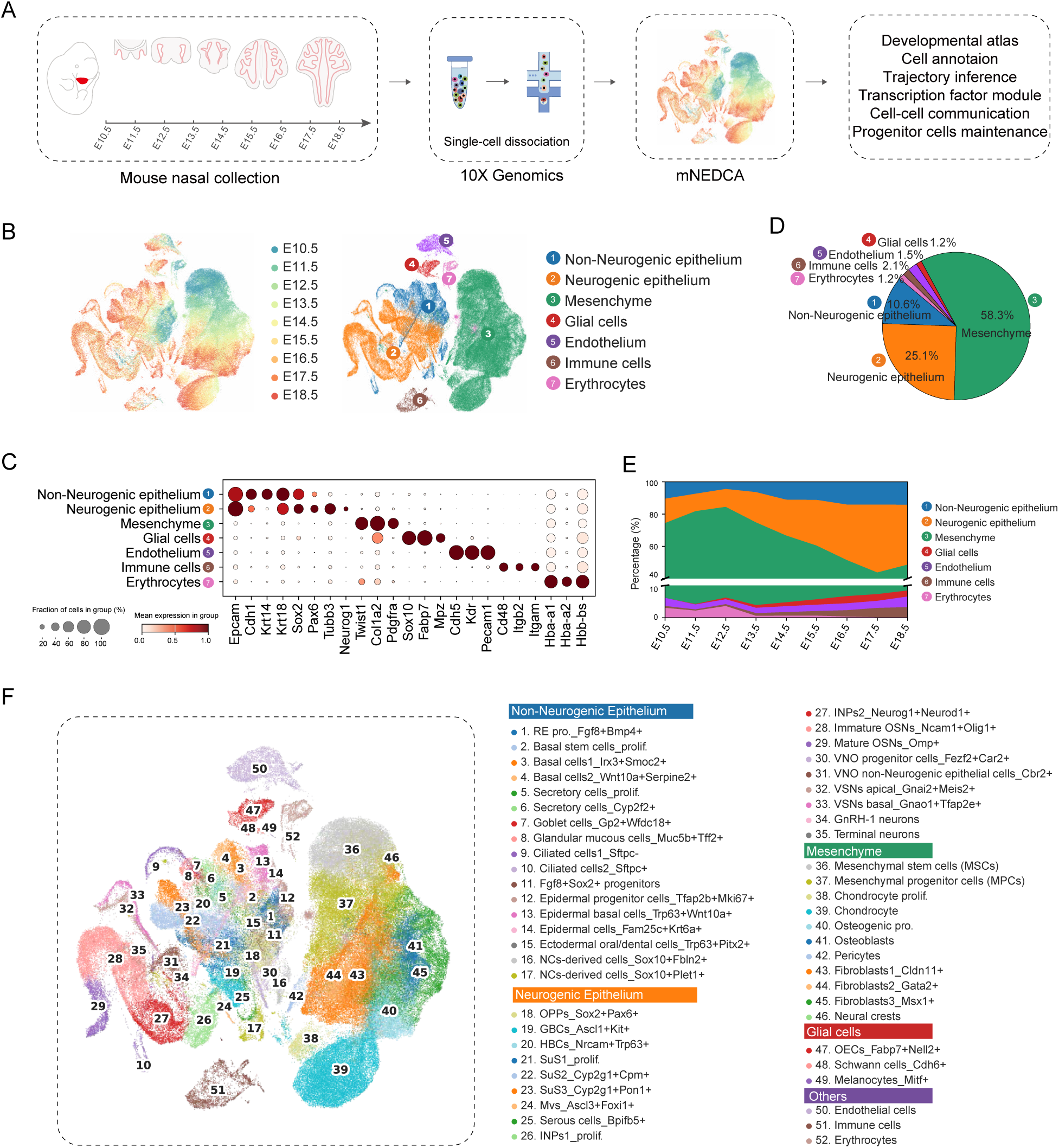
Single-cell atlas of mouse embryonic nasal development. A: Schematic overview of the scRNA-seq experimental design and analysis workflow for mouse nasal development from E10.5 to E18.5 (two biological replicates per time point). The resulting mouse nasal embryonic development cell atlas (mNEDCA) enables analysis of cell fate decisions, lineage trajectories, and intercellular communication. B: UMAP visualization of 183,247 cells from mouse embryonic nasal tissue samples (n_pcs = 40, n_neighbors = 10). Each dot represents a cell. Left, colored by embryonic stage; Right, colored by annotated major cell type. UMAP, Uniform Manifold Approximation and Projection. C: Dot plot showing expression of representative canonical markers for seven cell populations. Dot color indicates average expression level and dot size indicates the fraction of cells expressing each gene. D: Pie chart representing the global proportions of the seven cell populations during the nasal development. E: Stacked bar plot showing the fraction of seven cell populations across development stages (E10.5-E18.5). Colors correspond to panel (B). F: Overview of the 52 distinct cell clusters of the mNEDCA, defined by hierarchical annotation and iterative sub-clustering.

Utilizing this dataset, we first annotated and identified seven major cell populations (Fig. 1B and 1C). These included: non-neurogenic epithelium (*Epcam*^+^/*Cdh1*^+^/*Krt14*^+^) and neurogenic epithelium (*Sox2*^+^/*Pax6*^+^/*Tubb3*^+^) (Roskams et al., 1998; Amato et al., 2024; Taroc et al., 2025) which collectively expanded from 25.4% to 51.2% during development and formed the functional mucosal compartments of the nasal cavity (Figs. 1D, 1E, S1F). Mesenchymal cells (*Twist1*^+^/*Col1a2*^+^/*Pdgfra*^+^), constituted predominant population, decreasing from 58.3% to 40.2%, consistent with progressive epithelialization and structural remodeling (Fig. 1D and 1E). We also identified glial cells (*Sox10*^+^/*Fabp7*^+^/*Mpz*^+^), derived from neural crest lineages (Barraud et al., 2010; Forni et al., 2011; Katoh et al., 2011; Perera et al., 2020; Senf et al., 2025; Taroc et al., 2025), endothelial cells (*Pecam1*^+^/*Cdh5*^+^/*Kdr*^+^) that establish vascular networks, immune populations (*Cd48*^+^/*Itgb2*^+^/*Ptprc*^+^) and erythrocytes (*Hba-a1*^+^/*Hba-a2*^+^/*Hbb-bs*^+^), supporting metabolic supply (Fig. 1B–1D).

To further dissect the cellular heterogeneity within these major classes, we performed iterative sub-clustering and identified 52 molecularly distinct subtypes spanning progenitor, intermediate, and differentiated states across epithelial and mesenchymal lineages (Fig. 1F; Table S2). This analysis uncovered a spectrum of lineage-specific populations that emerged in a stage-dependent manner, providing a detailed view of nasal cellular diversification (Figs. 1F and S2A).

To delineate transcriptional dynamics related to developmental chronology, we performed correlation analysis of single-cell transcriptomes across all populations along the temporal continuum. The results revealed four transcriptionally distinct stages (Fig. S2B and S2C). Stage 1 (E10.5–E11.5) was characterized by enrichment of cell cycle regulation, telomere organization, and stem cell population maintenance, indicative of rapid proliferative expansion. Stage 2 (E12.5 – E13.5) featured gene programs associated with stem cell differentiation and epithelial morphogenesis, reflecting establishment of region-specific lineages. Stage 3 (E14.5–E15.5) showed progressive upregulation of extracellular matrix organization and cartilage development genes, consistent with formation of nasal structural frameworks (Kaucka et al., 2018). Stage 4 (E16.5–E18.5) was enriched with ATP metabolic process and metal ion and oxidative stress response, suggesting functional maturation of nasal tissue (Fig. S2B and S2C).

Collectively, we generated a high-resolution, temporally resolved mouse nasal embryonic development cell atlas (mNEDCA). This resource enables us to further identify more novel molecular and cellular features and investigate how different cell populations coordinate embryonic nasal development.

### Molecular characterization of the embryonic nasal epithelium

The nasal epithelium constitutes the primary interface for respiration and olfaction and also contributes to neuroendocrine and reproductive functions (Chung et al., 2016). It comprises multiple regionally specialized domains, including the respiratory epithelium (RE), olfactory epithelium (OE), vomeronasal organ (VNO), squamous epithelium (SE), septal organ (Ma et al., 2003; Tian and Ma, 2008), Grueneberg ganglion (Fuss et al., 2005), and migratory neuronal populations such as GnRH-1 and terminal neurons (Parslow et al., 2024). Despite this anatomical complexity, transcriptional profiles and reliable markers for these region-specific epithelial cell types during embryogenesis remain incompletely defined at single-cell resolution.

To delineate epithelial diversity, we performed in-depth analysis of the neurogenic and non-neurogenic epithelial compartments within mNEDCA, comprising 65,360 cells. Unbiased clustering resolved seven epithelial lineages corresponding to RE, OE, VNO, SE, GnRH-1 neurons, terminal neurons and a neural crest-like epithelial population (Fig. 2A and 2B). All clusters expressed canonical nasal epithelial markers (*Epcam/Six3*) (Rodenburg et al., 2023), whereas each lineage displayed distinct regional signatures (Fig. 2C; Table S3). The RE was characterized by *Foxa1* (Besnard et al., 2004; Wan et al., 2005; Paranjapye et al., 2020; Kusumoto et al., 2025; Taroc et al., 2025) and maintained a relatively stable proportion across stages (Figs. S3A, S3C, 2E). We identified previously underappreciated RE-enriched genes including *Serpinb11*, *Reg3g* and *Lox* (Fig. 2C), and functional enrichment with pathways involved morphogenesis of a branching epithelium and WNT signaling (Fig. S3B).

**Fig. 2.**
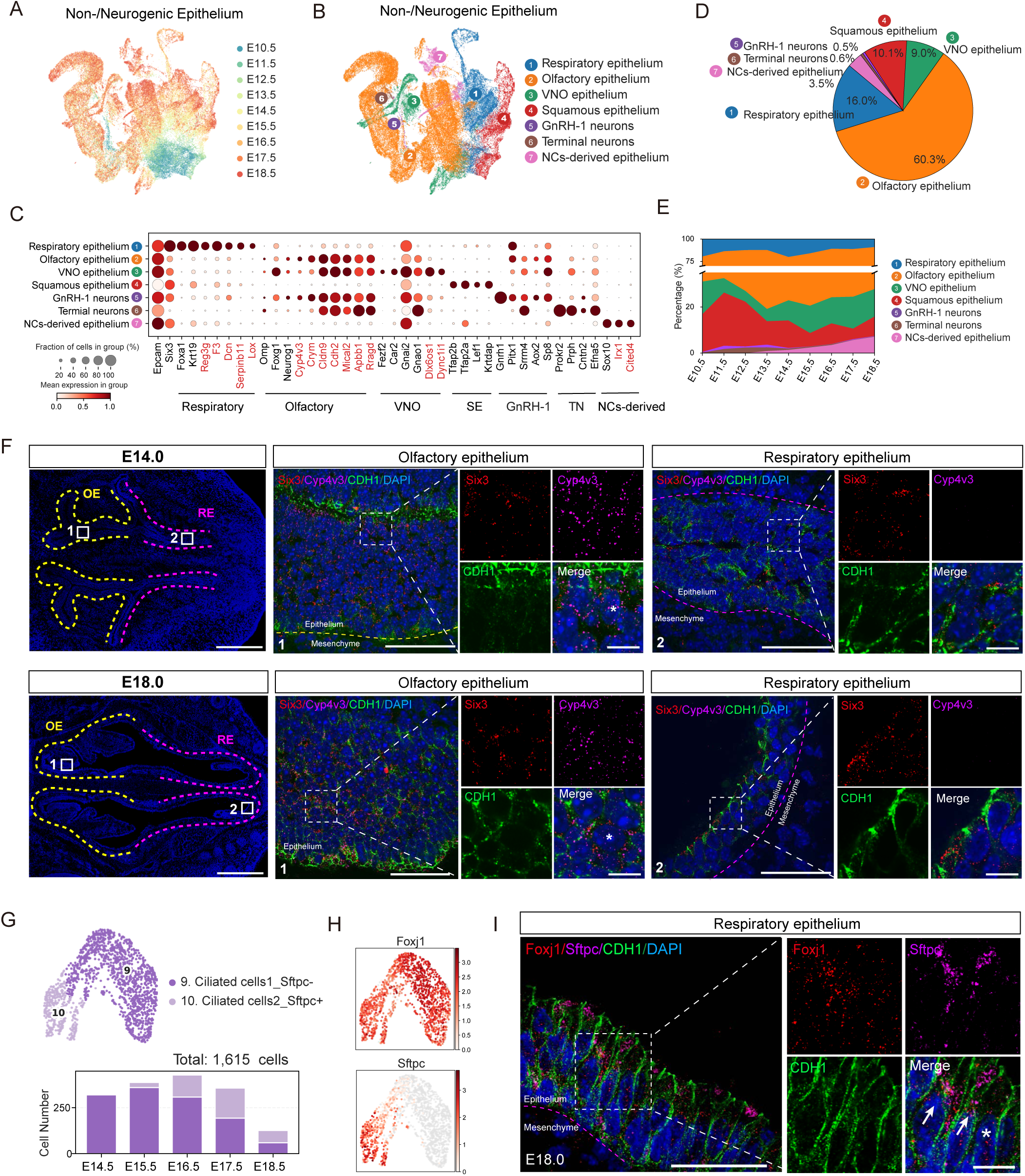
Molecular characterization of neurogenic and non-neurogenic nasal epithelial regions. A: UMAP visualization of 65,360 epithelial cells spanning neurogenic and non-neurogenic regions across developmental stages (n_pcs = 30, n_neighbors = 10). Each dot represents a cell. B: UMAP visualization of the subdivision of neurogenic and non-neurogenic epithelium into distinct regions (respiratory, olfactory, vomeronasal, squamous, GnRH-1 neurons, terminal neurons and NC-derived cells). Cells are colored by annotated region identity. C: Dot plot showing the expression of canonical markers and differentially expressed genes associated with different regions. Dot color indicates average expression level and dot size indicates the fraction of cells expressing each gene. Novel markers are highlighted in red. D: Pie chart summarizing the global proportions of several regions across all stages. Colors correspond to panel (B). E: Stacked bar plot illustrating the fraction of several regions across development stages. Colors correspond to panel (B). F: SmFISH combined with immunofluorescence staining of E14.0 (top) and E18.0 (bottom) nasal tissue. Low-magnification scans (left) provide an overview of respiratory and olfactory regions. High-magnification views showing specific *Cyp4v3* (magenta) expression restricted to olfactory epithelium (middle; asterisks denote individual positive cells), and absent from respiratory epithelium (right). *Six3* (red) marks nasal epithelium and CDH1 (green) immunofluorescence outlines epithelial boundaries. Images are representative of at least three independent experiments. Scale bars: 500 μm (left), 40 μm (middle/right); 10 μm for insets. G: UMAP visualization of 1,615 ciliated cells from the respiratory epithelium (BBKNN integration, n_pcs = 30, n_neighbors = 50). Accompanying bar plot shows the number of cells contributed by each developmental stage. H: UMAP feature plots showing the expression of *Foxj1* and *Sftpc* in the ciliated cells (color indicates normalized expression). I: SmFISH combined with immunofluorescence staining of *Foxj1* and *Sftpc* in E18.0 nasal respiratory epithelium. High-magnification views showing *Foxj1*^+^*Sftpc*^+^ ciliated cells (arrowheads) and *Foxj1*^+^*Sftpc*^-^ ciliated cells (asterisk). CDH1 (green) immunofluorescence labels epithelial boundaries. Images are representative of at least three independent experiments. Scale bars: 50 μm; 20 μm for insets.

The OE represented the largest epithelial population (Fig. 2D) and was defined by established markers *Omp* (Buiakova et al., 1996), *Foxg1* (Duggan et al., 2008), and *Neurog1* (Shaker et al., 2012). Beyond these classic signatures, our analysis highlighted *Cyp4v3*, *Cldn9* and *Rprm* as regional markers of OE (Fig. 2D). OE-enriched genes were associated with axonogenesis and synapse assembly (Fig. S3B). smFISH confirmed OE-specific expression of *Cyp4v3* (Fig. 2F), a member of the cytochrome P450 family, which participated in lipid metabolism and steroid hormone biosynthesis (Jia et al., 2023). *Cldn9*, encoding a tight junction protein, may contribute to olfactory epithelium barrier integrity (Nakano et al., 2009; Bird et al., 2010).

Apart from RE and OE, the rodent-specific structure vomeronasal organ (VNO) was well distinguished by typical markers *Fezf2*/*Car2* (Eckler et al., 2011) and novel specific genes like *Dlx6os1* and *Dync1i1*. VNO associated genes were enriched in cellular response to monoamine stimulus and hormone secretion (Fig. S3B). Squamous epithelium (SE) in the nasal vestibule expressed canonical keratinocyte markers *Krtdap*/*Tfap2b* (Van Otterloo et al., 2022) and was characterized by enrichment of skin development and keratinocyte differentiation (Fig. S3B). We also identified *Gnrh1*^+^/*Axo2*^+^ GnRH-1 neurons and *Prokr2*^+^/*Prph*^+^ terminal neurons (Figs. 2C and S3A) (Duittoz et al., 2022; Amato et al., 2024). Unlike other epithelial clusters above appearing at E10.5, the neural crest-derived epithelial subset emerged at E13.5 and specifically express *Sox10*/*Foxc1* as well as novel genes like *Irx1*/*Cited4*, showing robust enrichment in neural crest cell development, differentiation and migration (Figs. 2C and S3B).

Among these major epithelial lineages, embryonic RE heterogeneity has been poorly characterized. Sub-clustering resolved ten RE populations with distinct molecular identities (Fig. S4A and S4B). Progenitors were marked by *Fgf8*/*Dlk1*/*Bmp4* (Kawauchi et al., 2005; Maier et al., 2010; Kitagaki et al., 2011; Maier et al., 2011; Forni et al., 2013), whereas the basal compartment (*Trp63*⁺/*Dlk2*⁺) comprised a proliferative *Top2a*⁺ subset (Fig. S4B) (Deprez et al., 2020) and two differentiating branches defined by *Irx3*⁺/*Smoc2*⁺ and *Wnt10a*⁺/*Serpinb2*⁺ (Fig. S4B). Secretory lineages (*Cyp2f2*⁺/*Serpinb11*⁺) segregated into proliferative secretory cells, mature secretory cells, *Gp2*⁺*Wfdc18*⁺ goblet cells, and *Muc5b*⁺*Tff2*⁺ glandular mucous cells (Fig. S4B) (Ualiyeva et al., 2024). Strikingly, within the ciliated population (*Foxj1*⁺/*Deup1*⁺/*Myb*⁺) (Yu et al., 2008), we identified a previously unrecognized subcluster specifically expressing *Sftpc* (Fig. 2G and 2H), a canonical alveolar type II marker (Treutlein et al., 2014). smFISH and immunofluorescence confirmed co-localization of *Sftpc* with *Foxj1* and acetylated tubulin (Figs. 2I and S4C). *Sftpc*⁺ ciliated cells were enriched for genes involved in microtubule-based motility and cilium movement, whereas *Sftpc*⁻ ciliated cells preferentially expressed cell-cycle and ciliogenesis regulators (*Cdkn1a*, *Stmn1*, *Ccno*) (Fig. S4D and S4E; Table S4). These findings suggest functional diversification within RE ciliated cells and implicate the *Sftpc*⁺ subset in surfactant-related regulation of nasal mucosal homeostasis.

We next dissected OE heterogeneity and annotated 12 populations (Fig. S5A–S5C). Stem and progenitor compartments included *Sox2*⁺/*Pax6*⁺/*Rprm*⁺ olfactory progenitors (Krolewski et al., 2012), *Sox2*⁺/*Ascl1*⁺/*Kit*⁺ globose basal cells (Paronett et al., 2023), and *Trp63*⁺/*Krt14*⁺/*Nrcam*⁺ horizontal basal cells (Kawauchi et al., 2004; Packard et al., 2011). Neurogenic lineages comprised proliferative *Neurod1*⁺/*Mki67*⁺ immediate neuronal precursors (INPs), *Neurog1*⁺ INPs, *Ncam1*⁺ immature olfactory sensory neurons (OSNs), and Omp⁺ mature OSNs (Fig. S5B) (Graziadei et al., 1980; Cau et al., 2002; McIntyre et al., 2010; Ualiyeva et al., 2024). Remarkably, INPs and immature OSNs were detectable as early as E10.5, coinciding with the emergence of migratory GnRH-1 and terminal neurons (Figs. S5C and 2E), consistent with rapid initiation of olfactory neurogenesis following pit formation (Cau et al., 2002; Taroc et al., 2020). Non-neurogenic lineages included sustentacular trajectories from proliferative *Cyp2g1*⁺/*Mki67*⁺ progenitors (Gu et al., 1999) to mature *Ermn*⁺/*Cpm*⁺ and *Cyp2g1*⁺/*Pon1*⁺ states, as well as *Ascl3*⁺/*Foxi1*⁺ microvillar and *Bpifb5*⁺/*Kcnn4*⁺ serous cells (Fig. S5B) (Ualiyeva et al., 2024). Comparable subtype diversity was also resolved in VNO (Katreddi and Forni, 2021; Katreddi et al., 2022; Lin et al., 2022; Hills et al., 2024; Dietz et al., 2025), SE (Yu et al., 2020; Ye et al., 2022), and neural crest–derived compartments (Fig. S6A–S6F).

Together, these analyses define the molecular landscape of embryonic nasal epithelial lineages and reveal regional and intra-lineage heterogeneity. In particular, the discovery of a *Sftpc*⁺ ciliated population highlights previously unappreciated functional diversity within the respiratory epithelium and provides new entry points for investigating nasal epithelial specialization.

### Heterogeneity of the embryonic nasal mesenchyme

In addition to the nasal epithelium, the nasal mesenchyme plays an indispensable role in nose development. Mesenchyme-derived cavity structures define nasal shape and, together with the soft palate and mouth, contribute to voice production (LaMantia et al., 2000; Yang et al., 2018). In our mNEDCA dataset, nasal mesenchyme represented the largest population among seven major cell clusters, underscoring its essential function in providing the structural matrix for nasal morphogenesis. However, the precise cellular composition and dynamic changes of this compartment during embryonic development remain poorly understood.

To this end, we performed an in-depth analysis of 105,140 mesenchymal cells, classifying them into ten transcriptionally distinct subpopulations using established marker genes (Fig. 3A–3C). Among them, two progenitor populations were identified: mesenchymal stem cells (MSCs) and mesenchymal progenitor cells (MPCs). Both exhibited high expression of stemness markers *Pax7*/*Hmga2* (Murdoch et al., 2010) and proliferation-associated genes (*Top2a*, *Mki67*) but showed minimal expression of lineage-specific differentiation markers (Fig. 3C). Temporally, MSCs predominated at early developmental stages but disappeared after E12.5, whereas MPCs emerged at E11.5 and persisted until E18.5 (Fig. 3D and 3E). While MSCs and MPCs shared numerous cell cycle and stemness related genes, they displayed distinct transcriptional programs. MSCs were enriched for genes regulating serine/threonine kinase activity and stem cell maintenance, whereas MPCs expressed genes involved in extracellular matrix organization, connective tissue development, and cell fate specification (Fig. 3F and 3G). These observations suggest MPCs serve as a fate-committed intermediate between MSCs and differentiated mesenchymal lineages.

**Fig. 3.**
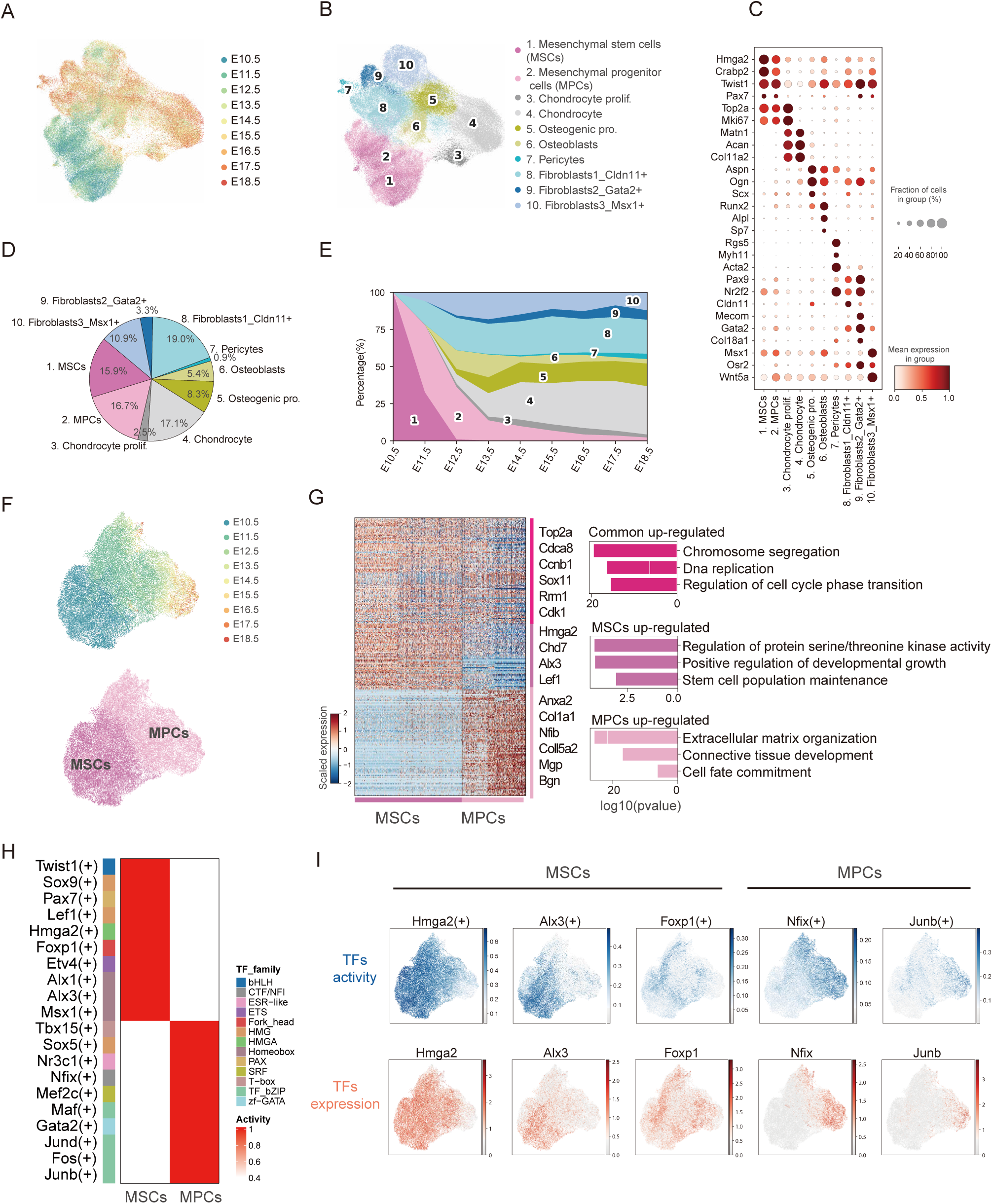
Classification of nasal mesenchymal cells and key transcriptional regulation of MPCs. A: UMAP visualization of 105,140 nasal mesenchymal cells colored by different embryonic stages (Harmony integration, n_pcs = 30). Each dot represents a cell. B: UMAP projection showing annotated major mesenchymal populations. C: Dot plot showing the expression of marker genes across mesenchymal populations. Dot color indicates average expression level and dot size indicates the fraction of cells expressing each gene. D: Pie chart representing the global proportions of major mesenchymal cell types. Colors correspond to panel (B). E: Stacked bar plot illustrating the fraction of each major mesenchymal cell type across embryonic stages. Colors correspond to panel (B). F: UMAP visualization of Mesenchymal stem cells (MSCs) and Mesenchymal progenitor cells (MPCs) colored by embryonic stages (BBKNN integration, n_pcs = 30). Each dot represents one cell. G: Left, heatmap of genes shared or differentially expressed between MSCs and MPCs (Wilcoxon rank-sum test, *P* < 0.01), selected genes are labelled. Right, GO enrichment analysis of the identified DEGs, significance is indicated by *P*-values. H: Heatmap of regulon activity (AUC scores) in MSCs versus MPCs inferred by pySCENIC. AUC scores were calculated at the single-cell level and averaged per cell type. Regulons are row-z-score normalized. TF families are annotated on the left. Regulons were filtered for measurable activity (see Methods). I: UMAP feature plots showing the TF activity (blue, AUC score) and gene expression (red, log-normalized) of MSCs-specific (*Hmga2*, *Alx3* and *Foxp1*), and MPCs-specific (*Nfix* and *Junb*). SCENIC-generated TF activity, represented by AUC score, reflects the co-expression strength of TF and its target genes.

The remaining eight mesenchymal subtypes were classified into four main differentiation lineages: chondrogenic, osteogenic, fibroblastic, and pericytic. Chondrocytes (*Matn1*⁺/*Acan*⁺) (Hojo and Ohba, 2019) secreted type II collagen and proteoglycans, while osteoblasts (*Aspn*⁺/*Ogn*⁺/*Runx2*⁺) (Fujita et al., 2004) formed nasal bones and cribriform plates, providing mechanical support. Pericytes (*Rgs5*⁺/*Acta2*⁺) (Berger et al., 2005) contributed to microvascular stability, and fibroblasts mediated connective tissue morphogenesis and basement membrane scaffolding (Figs. 3C and S7A; Table S5). Consistent with the stage 3 developmental features above (Fig. S3C and S3D), these differentiated cell types expanded markedly after E13.5 (Fig. 3E), indicating their crucial role in establishing nasal architecture.

Next, to verify MPCs as potential fate-committed intermediate, we employed RNA velocity and partition-based graph abstraction (PAGA) analysis (Wolf et al., 2019) to reconstruct lineage trajectories of nasal mesenchyme. As expected, the result revealed that MSCs transition through MPCs before differentiating into multiple mesenchymal lineages (Fig. S7B–S7D). Following this developmental trajectory, we performed single-cell regulatory network inference (SCENIC) analysis (Aibar et al., 2017) to investigate key transcription factors (TFs) governing nasal mesenchymal cell fate transition. Consistent with previous reports, the identification of known TFs like *Hmga2* and *Alx3* (Fig. 3H and 3I) (Beverdam and Meijlink, 2001; Negishi et al., 2022) which highly expressed and activated in MSCs validate the analytical approach. Furthermore, we showed novel TFs regulators such as *Foxp1*/*Etv4* in MSCs and *Nfix*/*Junb* in MPCs, implying their regulatory roles in fate determination (Fig. 3H and 3I).

On the other hand, the subsequent differentiation programs were orchestrated by distinct lineage-determining transcriptional regulators. For instance, *Sox9* and *Plagl1* governed chondrogenesis (Akiyama, 2008), regulating both early commitment and terminal maturation. *Maf* and *Klf13* emerged as regulator directing fibroblast lineage specification (Fig. S7E and S7F). The osteogenic lineage was controlled by a *Dlx5* that progressively upregulated bone matrix genes (Samee et al., 2008), while *Ets1* and *Mef2c* regulated pericyte differentiation (Fig. S7E and S7F). Several additional TF candidates were identified that may represent novel regulators requiring further investigation.

Collectively, our results revealed the transcriptional diversity of nasal mesenchyme and support a model in which MPCs function as fate-committed intermediates guiding mesenchymal cell lineage differentiation during nasal morphogenesis.

### Diverse signals governing RE progenitor cells maintenance

The rich diversity and distinct origins of cells participating in embryonic nasal development imply intricate intercellular communication underlying nasal morphogenesis. To elucidate these interactions, we conducted systematic ligand–receptor analyses across major cell types in mouse nasal development. Quantitative mapping of outgoing and incoming signals identified the nasal non-/neurogenic epithelium, mesenchyme, and glial cells as predominant communicators (Fig. 4A and 4B), with mesenchymal (e.g., *Bmp4*, *Wnt5a*) and glia-derived signals (e.g., *Wnt6*, *Dhh*, *Bmp7*) collectively accounting for ∼50% of epithelial inputs, highlighting the established roles of BMP, WNT, and NOTCH pathways in nasal development (LaMantia et al., 2000; Sokpor et al., 2018). The segregation of olfactory and respiratory epithelium is directed by antagonistic FGF and BMP signaling (Maier et al., 2010). In both chick and mouse models, elevated BMP activity, marked by pSmad1/5/8 and activation of *Msx1/2* and *Id3*, promotes respiratory epithelial fate, whereas BMP antagonists, including *Noggin*, are required to preserve the neurogenic sensory domain (Forni et al., 2013). In addition, autocrine signals from the epithelium (*Fgf9*, *Nrg1*, *Ocln*) also contributed to self-regulation (Fig. S8A). These results suggested that paracrine and autocrine signals cooperatively modulated global nasal epithelial development.

**Fig. 4.**
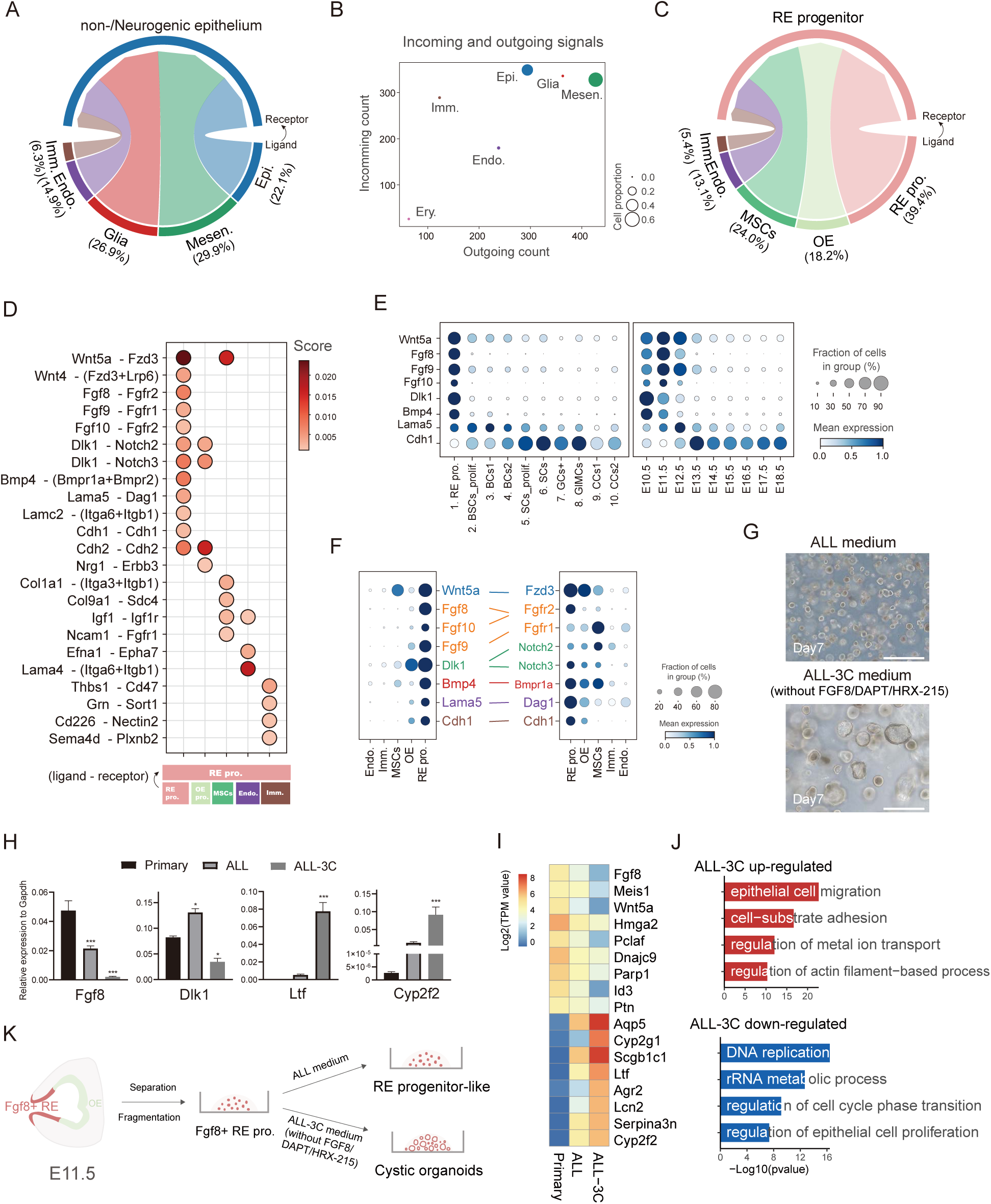
Niche homeostasis signaling in early respiratory epithelial progenitor cells. A: Circos plot of ligand-receptor (LR) interactions between non-/neurogenic epithelium and other major cell types across E10.5-E18.5. Ribbon widths represent the interaction weight. LR inference was performed with CellChat (see Methods). Epi., non-/neurogenic epithelium; Mesen., nasal mesenchyme; Glia, glia cells; Endo., endothelium; Imm., immune cells. B: Dot plot showing the outgoing (secreted) and incoming (received) signals counts among major cell types. Dot size indicates population proportion. C: Circos plot of LR interactions between RE progenitor cells and other cell types across E10.5-E11.5. Ribbon widths represent the interaction weight. RE pro., RE progenitor cells; OE, olfactory epithelium; MSCs, Mesenchymal stem cells; Endo., Endothelium; Imm., Immune cells. D: Dot plot showing LR interaction strength related to panel (C). The color of each dot represents the communication probability. Only significant LR pairs are displayed (filtering criteria detailed in Methods). E: Dot plot showing the expression of ligands in RE pro. across RE cell types (left) and developmental stages (right). BSCs, basal stem cells; BC, basal cells; SC, secretory cells; GC, goblet cells; GMC, glandular mucous cells; CC, ciliated cells. F: Paired dot plots presenting the expression of ligands (left) and receptors (right) among RE pro. and interacting populations, color-coded by LR pairs. Dot color indicates average expression level and dot size indicates the fraction of cells expressing each gene. G: Representative morphologies images of RE progenitor cells cultured after 7 days of induction in ALL and ALL-3C (without FGF8, DAPT and HRX215) medium. Images are representative of at least three independent experiments. Scale bars, 100 μm. H: qRT–PCR quantification of progenitor marker (*Fgf8*, *Dlk1*) and differentiation marker (*Ltf* and *Cyp2f2*) after 7 days in ALL or ALL-3C medium. Data are mean ± SEM from *n* = 3 independent biological replicates. Statistical comparison was performed by ordinary one-way ANOVA followed by Dunnett’s post-hoc test; significance thresholds are indicated in the panel (* *P* < 0.05; ** *P* < 0.01; \*\*\**P* < 0.001 vs. the Primary RE progenitor cells). I: Heatmap of RE progenitor-related and differentiation-related genes after 7 days in ALL or ALL-3C medium. Gene expression values were log-normalized for visualization. J: Bar plot displays the GO terms enriched among genes that are upregulated or downregulated in ALL-3C compared to ALL medium, with significance indicated by *P*-values. K: Schematic model of in vitro maintenance of RE progenitor cells. Cells form compact spheroid in ALL medium but differentiate into cystic structures upon removal of FGF8, DAPT, and HRX215 (All-3C medium).

Focusing on epithelial subtypes, we found that OE development was primarily driven by mesenchymal inputs (32.5%), with WNT pathways critically regulating sensory cell fate and morphogenesis, which resembled global epithelial cells communication pattern (Kawauchi et al., 2004) (Fig. S8B and S8C). However, RE development relied more heavily on autocrine signaling (34.0%) than that from mesenchyme, glia or other cell types (Fig. S8D and S8E). This suggested that RE development might involve special signaling patterns, which inspired us to conduct further analysis of RE subclusters.

Then, we performed ligand-receptor interaction analysis for respiratory epithelial progenitor cells (RE pro.), whose microenvironments for self-renewal and differentiation were previously underestimated due to unavailable in vitro culture models. The results revealed that autocrine signaling accounted for the highest proportion with 39.4% (Fig. 4C). Specifically, it involves autocrine signaling pathway including NOTCH (*Dlk1*-*Notch2/3*), FGF (*Fgf8*/*Fgf10*-*Fgfr2* and *Fgf9*-*Fgfr1*) and WNT (*Wnt5a*-*Fzd3*) as well as ECM signaling (*Col9a1*-*Sdc4*) (Fig. 4D; Table S6). Furthermore, we observed that ligands *Dlk1*, *Fgf8*, *Fgf10* and *Wnt5a* highly expressed in RE pro. during early development, but declined at later stages (Fig. 4E and 4F), suggesting a role in progenitor homeostasis.

To test this, we decided to utilize these signals agonist, inhibitor and their combinations to establish an in vitro culture system of RE progenitor cells isolated from E11.5 nasal epithelial tissues. Ultimately, we developed one chemically defined medium (“All” medium) to capture RE progenitors in vitro, which consisted of three core small molecules FGF8, DAPT (NOTCH inhibitor) and HRX215 (WNT/NOTCH modulator) (Zwirner et al., 2024) alongside supplementary factors. Under these conditions, RE progenitor cells maintained compact and smooth spheroid morphology (Fig. 4G), and sustained the expression of progenitor markers *Fgf8*, *Dlk1*, *Wnt5a* and *Hmga2* by both qPCR and bulk RNA-seq (Fig. 4H and 4I). On the contrary, removal of the three core factors (“All-3C” medium) induced cystic structures from compact morphology rapidly (Fig. 4G). Furthermore, we observed that differentiated genes such as *Aqp5*, *Cyp2g1* and *Scgb1c1* upregulated obviously in the absence of these three core small molecules (Fig. 4I). GO term analysis showed upregulated genes were enriched in epithelial cell migration and regulation of metal ion transport while decreased genes were related to DNA replication, rRNA metabolic process and cell cycle regulation (Fig. 4J). These results confirmed that the FGF, WNT and NOTCH signaling played important roles in establishing essential microenvironments for RE progenitor cell maintenance in vitro (Fig. 4K).

Together, these analyses reveal a previously unappreciated autocrine-dominant niche governing RE progenitors and provide evidence that FGF/NOTCH/WNT mediated circuits sustain progenitor identity. The culture system established here offers a platform for mechanistic dissection of respiratory epithelial lineage specification.

### Foxa1 as a key regulator of respiratory epithelium differentiation

The in vitro culture system for RE progenitor cells described above enabled further investigation of key regulators of RE lineage differentiation predicted by the mNEDCA dataset. Among these, transcription factors (TFs) represent major drivers of cell fate transitions during development. To elucidate the roles of TFs in RE development, we first performed trajectory analysis using RNA velocity (Figs. 5A and S9A), which revealed that RE progenitor cells serve as the developmental origin, giving rise to proliferative basal stem cells. These basal stem cells subsequently diverged into two distinct lineages: a basal cell lineage generating *Irx3*⁺ and *Wnt10a*⁺ basal cell subtypes, and a secretory–ciliated lineage in which secretory progenitors either differentiated into secretory cells or *Gp2*^+^*Wfdc18*⁺ goblet cells or underwent terminal differentiation into *Foxj1*⁺*Myb*⁺ ciliated cells (Fig. 5A–5C, S9B). This bifurcating differentiation program was consistently supported by trajectory inference analyses with Monocle2 (Qiu et al., 2017) (Fig. S9C).

**Fig. 5.**
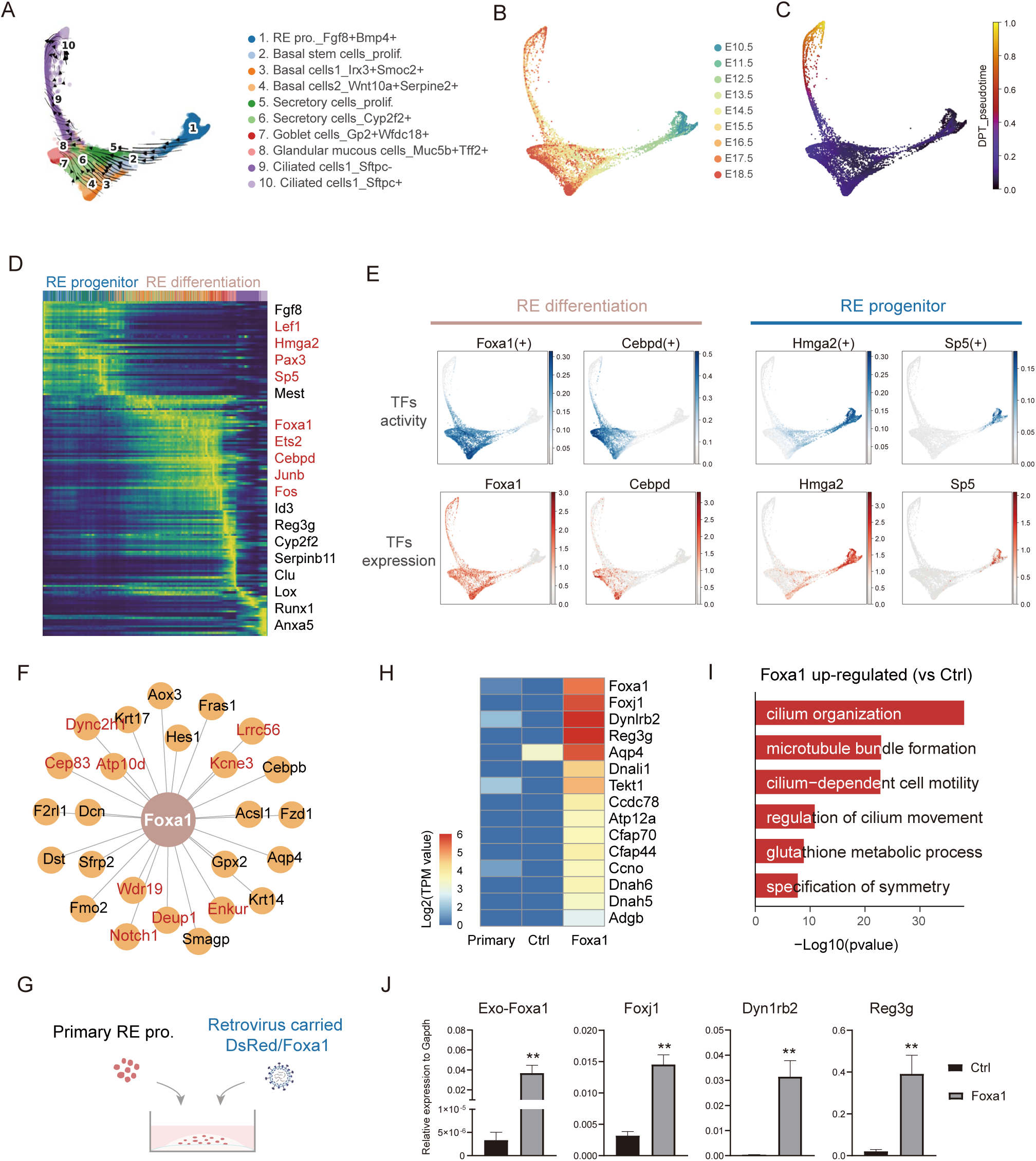
Foxa1 as a key regulator of nasal respiratory epithelium differentiation. A: PHATE visualization of the respiratory epithelium colored by cell types (10,432 cells). RNA velocity vectors were projected onto the graph to infer directional lineage progression (see Methods). PHATE, Potential of Heat-diffusion for Affinity-based Trajectory Embedding. B: PHATE embedding colored by embryonic stage, showing the temporal distribution of respiratory epithelial cells. C: PHATE graph of respiratory epithelium colored by Diffusion Pseudotime (DPT). The gradient from dark purple to bright yellow represents the predicted differentiation progression. D: Heatmap of genes differentially expressed along pseudotime trajectory; selected genes are labelled and TFs are highlighted in red (Wilcoxon rank-sum test, *P* < 0.01). E: PHATE graph showing the TF activity (blue, AUC score) and gene expression (red, log-normalized) for RE differentiation-specific TFs (*Foxa1* and *Cebpd*), and RE progenitors-specific TFs (*Hmga2* and *Sp5*). TF activity was inferred using pySCENIC. F: Transcriptional regulatory network centered on *Foxa1* nodes; edges denote regulatory links retained after cis-motif pruning (pySCENIC). Genes associated with ciliary function are highlighted in red. G: Schematic of the in vitro overexpression experiment in primary RE progenitor cells. Cells were transduced with *Foxa1* overexpression (Foxa1) or DsRed (Ctrl), and cultured under defined induction conditions in ALL medium. H: Heatmap showing the differential gene expression between Ctrl (DsRed) and Foxa1 overexpression conditions. Gene expression values (TPM values) were log-normalized for visualization. I: Bar plot displaying the GO terms enriched among genes that are upregulated in *Foxa1* overexpression compared to Ctrl (DsRed), with enrichment significance indicated by *P*-values. J: qRT-PCR quantification of Exo-*Foxa1*, *Foxj1* and *Sftpc* expression in Foxa1 overexpression versus Ctrl (DsRed). Data are mean ± SEM from n = 3 independent biological replicates. Statistical comparison was performed by two-tailed unpaired *t*-test; significance thresholds are indicated in the panel (* *P* < 0.05; ** *P* < 0.01 vs. the Ctrl).

To identify TFs initiating RE differentiation, we performed comparative transcriptomic analysis between RE progenitor cells and their differentiated counterparts (RE differentiated cells) (Fig. S9D). RE progenitor cells exhibited high expression of stemness-associated genes (*Fgf8*) and TFs such as *Hmga2*, *Pax3*, and *Sp5*. In contrast, differentiated cells showed progressive upregulation of maturation markers (*Reg3g*, *Cyp2f2*, *Anxa5*) as well as TFs including *Foxa1*, *Cebpd*, and *Junb* along pseudotemporal ordering (Figs. 5D and S9E; Table S7). SCENIC analysis of TF–gene regulon activity further identified *Hmga2* and *Sp5* as key regulators maintaining RE progenitor cell identity, while *Foxa1* and *Cebpd* emerged as central drivers orchestrating RE lineage differentiation (Fig. 5E). Notably, SCENIC predicted *Deup1*, *Kcne3*, *Atp10d*, *Wdr19*, and *Aqp4* as downstream targets of *Foxa1*, all of which are activated during differentiation and participate in cilium assembly and motility pathways (Figs. 5F and S9F). Consistent with these predictions, smFISH confirmed restricted *Foxa1* expression within the RE region (Fig. S3C).

To functionally validate *Foxa1*, we overexpressed Foxa1 in RE progenitor cells via retroviral transduction (Figs. 5G and S9G), which significantly upregulated ciliogenesis-related genes including *Foxj1*, *Dynlrb2*, *Aqp4*, and *Ccno* as revealed by RNA-seq (Fig. 5H). Gene ontology (GO) analysis indicated enrichment for pathways related to cilium movement and glutathione metabolism (Fig. 5I), which was further confirmed by RT–qPCR validation (Fig. 5J).

Together, these findings delineated the differentiation trajectory of the respiratory epithelium and demonstrated that *Foxa1* functions as a key regulator of RE differentiation, directing progenitor cells toward the ciliated cell lineage.

### Olfactory epithelial stem cell dynamics

Compared with respiratory epithelial progenitor cells, olfactory epithelial (OE) stem cells have attracted greater attention, yet the precise identity of stem cell populations within the OE remains under debate. Globose basal cells (GBCs) have been proposed as one candidate, given their molecular features and phenotypic similarity to olfactory placode progenitor cells (OPPs), which generate the OE during embryogenesis (Kawauchi et al., 2004; Packard et al., 2011; Krolewski et al., 2012). However, accumulating evidence indicates that GBCs constitute a heterogeneous population (Krolewski et al., 2013; Schwob, J. E. et al., 2017). This apparent paradox prompted us to investigate OE stem cell dynamics during nasal development.

We re-clustered the OPPs and GBCs characterized above and found that they segregated into two distinct clusters (Fig. 6A). One cluster represented OPPs, which peaked at embryonic day 10.5 (E10.5) and rapidly declined after E12.5, whereas the other cluster comprised GBCs, which emerged at E12.5 and persisted until E18.5 (Fig. 6B). This temporal transition aligns with the established model of olfactory neurogenesis (Packard et al., 2011; Kim et al., 2023), in which GBCs arise prior to horizontal basal cells (HBCs) and represent predominant stem cell pool at embryonic stages. Both two clusters maintained an undifferentiated state with expression of canonical olfactory stemness markers *Sox2* and *Pax6* (Schwob, J. E. et al., 2017), and shared molecular features related to DNA replication and cell cycle regulation (Fig. 6C and 6D). However, GBCs exhibited high expression of late-stage olfactory stemness markers *Ascl1* and *Kit*, which were minimally expressed in OPPs (Fig. 6C). Differential gene expression (DGE) analysis revealed striking transcriptional differences between OPPs and GBCs (Fig. 6D). OPPs specifically expressed a set of previously uncharacterized genes, including *Sall4*, *Mest*, *Igf2bp1*, and *Car14* (Fig. S10A). Notably, *Car14*, whose deficiency has been linked to liver fibrosis (Qian et al., 2022), showed specific co-localization with *Sox2* in the E11.5 OE by smFISH (Fig. 6E), suggesting a role in olfactory stem cell regulation. In contrast, GBCs specifically expressed *Lgr5*, a known adult GBC marker (Chen et al., 2014), along with other uncharacterized genes such as *Tmprss4*, *Jag2* and *Runx1* (Fig. S10A). *Lgr5* expression in *Ascl1*⁺ GBCs at E14.0 was further validated by smFISH (Fig. 6E). Gene ontology (GO) analysis revealed that OPPs-enriched genes were associated with epithelial cell proliferation and Wnt signaling, whereas GBCs-enriched genes were linked to Notch signaling and glial cell differentiation (Fig. 6D). SCENIC analysis identified distinct regulons, including *Tcf7l1* in OPPs and *Tcf4* in GBCs (Fig. S10B and S10C).

**Fig. 6.**
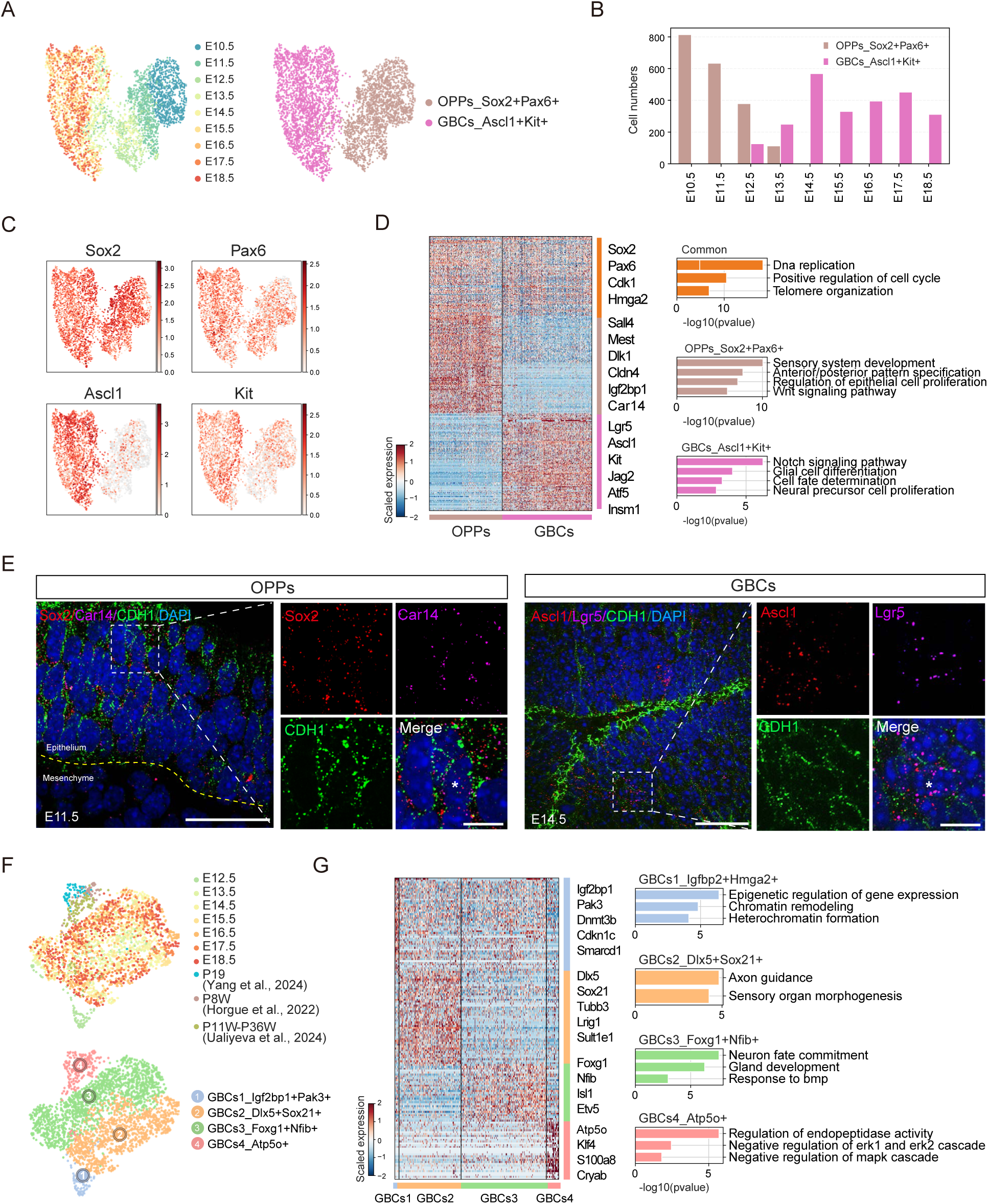
Differential characteristics of OPPs and GBCs and the heterogeneity during GBCs development. A: UMAP visualization of OPPs and GBCs (5,112 cells) colored by cell type (left) and embryonic stage (right) (n_pcs = 30, n_neighbors = 10). Each dot represents a cell. B: Bar plot indicating the cell counts of OPPs and GBCs across developmental stages. C: UMAP showing the expression of canonical marker genes *Sox2*, *Pax6*, *Ascl1* and *Kit* (log-normalized). *Sox2* and *Pax6* mark both OPPs and GBCs, while *Ascl1* and *Kit* are enriched in GBCs and low in OPPs. D: Left, heatmap of genes shared or differentially expressed between OPPs and GBCs (Wilcoxon rank-sum test, *P* < 0.01), selected genes are labelled. Right, GO enrichment analysis of the identified DEGs, significance is indicated by *P*-values. E: SmFISH and immunofluorescence staining of E11.5 (left) and E14.5 (right) nasal olfactory epithelium. High-magnification views showing specific *Sox2* (red) and *Car14* (magenta) expression restricted to OPPs (Left; asterisks denote individual positive cells), and specific *Ascl1* (red) and *Lgr5* (magenta) expression restricted to GBCs (right; asterisks denote individual positive cells). CDH1 (green) immunofluorescence outlines epithelial boundaries. Images are representative of at least three independent experiments. Scale bars, 40 μm; 10 μm for insets. F: UMAP visualization of GBCs (2,714 cells) across different embryonic and postnatal stages (top) and subdivided into four subtypes (bottom) (n_pcs = 30, n_neighbors = 15). P19: Postnatal 19 day (Yang et al., 2024); P8W: Postnatal week 8 (Horgue et al., 2022); P11W-P36W: Postnatal week 11 to 36 (Ualiyeva et al., 2024). G: Left, heatmap of differentially expressed genes among the four GBC subtypes (Wilcoxon rank-sum test, *P* value < 0.01), selected genes are labelled. Right, bar plot displays GO terms enriched among these genes, with enrichment significance indicated by *P*-values.

To further dissect heterogeneity of GBCs, we extended the single-cell timeline of nasal development by integrating our mNEDCA dataset with published postnatal datasets (Horgue et al., 2022; Ualiyeva et al., 2024; Yang et al., 2024) spanning P19 to adult (P36W). Based on transcriptional features, GBCs were classified into four subgroups with continuous developmental relationships (Fig. 6F). GBCs1, enriched for *Igf2bp1* and *Pak3* and associated with chromatin remodeling and heterochromatin formation, may represent transitional cells bridging OPPs and specialized GBCs (Fig. 6G and S10D). Prenatally, GBCs2 expressed *Dlx5* and *Sox21*, linked to axon guidance and sensory organ morphogenesis, whereas GBCs3 expressed *Foxg1* and *Nfib*, associated with neuronal fate commitment and gland development (Fig. 6G and S10D). These two clusters corresponded to the “active” GBC states described in adult neurogenesis (Schwob, James E et al., 2017). Postnatally, GBCs4, expressing *Atp5o* and *Klf4* and enriched for pathways including negative regulation of ERK1/2 signaling and modulation of endopeptidase activity, emerged as a mature population (Fig. 6G and S10D and Table S8). Pseudotime analysis confirmed a developmental trajectory from GBCs1 to active GBCs2/3, and ultimately to the homeostatic GBCs4 state (Fig. S10E–S10G).

Collectively, we delineate the dynamic transcriptional landscape of olfactory epithelial stem cells from E10.5 to adulthood, identifying novel markers that distinguish OPPs from GBCs during embryogenesis and refining our understanding of GBC heterogeneity and maturation.

## Discussion

In this study, we presented a high-resolution temporal single-cell RNA sequencing (scRNA-seq) atlas of mouse nasal development, profiling 183,247 cells from 20 embryos spanning embryonic day (E)10.5 to E18.5. We defined seven major cell classes and 52 subtypes, uncovered a series of previously unrecognized molecular markers, and identified rare or uncharacterized cell populations. Cell–cell communication analysis revealed critical niche-derived signals regulating respiratory epithelium (RE) progenitor cells maintenance, which guided the establishment of an in vitro RE progenitor cells culture system. Using this platform, we reconstructed the differentiation trajectory of RE lineages and demonstrated that Foxa1 functions as a key driver promoting nasal RE progenitors toward a ciliated epithelial fate. Furthermore, we delineated the transcriptional dynamics of olfactory epithelium (OE) stem cells from E10.5 through adulthood. The resulting mouse nasal embryonic development cell atlas (mNEDCA) offered a valuable reference for nasal development and regenerative medicine, particularly for stem cell-based strategies to restore nasal structure and function.

The nose serves as both a critical respiratory gateway and a sensory organ, and its development involves a complex interplay of diverse cell populations. The temporal resolution and deep transcript coverage of mNEDCA enabled systematic annotation of developmental stage–specific cell states. We performed unbiased clustering for each developmental stage, annotated clusters based on previously known markers and newly identified specific genes, which were validated them by smFISH or immunofluorescence in the context of anatomical structures. This approach uncovered novel markers, including *Cyp4v3* as an OE-enriched gene and *Car14* as an early OE stem cell associated marker. We also identified previously unreported cell types, such as *Sftpc*⁺ ciliated cell subset at late gestation and mesenchymal progenitor cells (MPCs) that serve as intermediates during nasal mesenchyme differentiation. Functional characterization of these genes and cell types will expand current understanding of nasal epithelial and mesenchymal development.

Among diverse cell populations, the epithelium plays a dominant functional role in nasal function and patterning. Analogous to other developing organs (Ribatti and Santoiemma, 2014; Le Guen et al., 2015; El Agha and Thannickal, 2023), nasal morphogenesis is shaped by reciprocal epithelial–mesenchymal interactions mediated by BMP, FGF, retinoic acid, and sonic hedgehog (SHH) signaling (LaMantia et al., 2000; Maier et al., 2010; Maier et al., 2011; Forni et al., 2013). Consistently, mNEDCA delineated multilayered inductive networks involving BMP, FGF, NOTCH, and WNT pathways across distinct cell populations. Notably, compared with other epithelial cell subtypes, RE development relies more heavily on its own autocrine signals. We identified FGF, NOTCH, and WNT circuits as core regulators of RE progenitor homeostasis and leveraged this insight to establish a chemically defined culture system. Within this context, we functionally validated *Foxa1* as a driver of RE differentiation toward the ciliated lineage. Although *Foxa1* expression in RE has been reported (Besnard et al., 2004), its developmental role was unclear. Our data indicate that *Foxa1* promotes ciliogenic programs, raising the possibility that *Foxa1* deficiency in vivo could impair mucociliary clearance and epithelial barrier integrity. Given the established role of *Foxa1* as a pioneer factor in lung epithelial development (Wan et al., 2005; Paranjapye et al., 2020), together with the identification of *Sftpc*⁺ ciliated subtypes in the nasal RE, suggests conserved mechanisms across the upper and lower respiratory tract. The RE progenitor system described here provides a promising platform for dissecting RE cell fate regulation and for isolating corresponding human stem cells for therapeutic applications in nose diseases.

In the OE, defining the precise stem cell populations remains an area of active debate. Globose basal cells (GBCs) and horizontal basal cells (HBCs) are two widely recognized candidates (Iwai et al., 2008; Fletcher et al., 2017). Our dataset captured both populations, as well as olfactory placode progenitors (OPPs) that give rise to diverse OE epithelial cell types. In agreement with previous studies (Packard et al., 2011; Kim et al., 2023), GBCs emerged earlier and were more abundant than HBCs, suggesting a potential primary role for GBCs in lineage specification during embryogenesis. We also resolved GBC heterogeneity across development and identified distinct markers separating GBCs from OPPs. Integration with postnatal datasets (Horgue et al., 2022; Ualiyeva et al., 2024; Yang et al., 2024) bridged embryonic and adult programs, although full reconstruction of OE lineages remains challenging due to plasticity and mixed origins. Definitive resolution will require lineage-tracing initiated at E9.5 or earlier and regionally targeted scRNA-seq from the frontonasal process.

Despite these findings, several limitations for mNEDCA should be considered. Cell coverage of mNEDCA at late developmental stages is low relative to the total nasal cellularity. This may lead to underrepresentation of rare cell types, such as Grueneberg ganglion which appear around E16.0 and contain less than 500 cells during embryogenesis (Fuss et al., 2005). Whole-tissue dissociation and droplet-based capture may bias against specialized neurons, including those of the septal organ. The absence of spatial transcriptomic information also limited discrimination of anatomically adjacent but transcriptionally similar populations like cells in septal organ, VNO and OE. Future integration of spatially resolved and region-specific profiling will refine the spatiotemporal architecture of nasal development. Moreover, although our analyses focused on epithelial and mesenchymal compartments, immune, endothelial, and neural crest–derived cells also contribute to morphogenesis and merit deeper exploration. Nevertheless, the mNEDCA provides a valuable resource to explore cellular heterogeneity, lineage dynamics, and regulatory networks during nasal embryogenesis, and offers novel insights into epithelial stem cell biology with implications for regenerative medicine.

## MATERIALS AND METHODS

### Animals

All animal experiments were performed in accordance with the Guide for the Care and Use of Animals Protocol (AUCP, GZLAB-AUCP-2024-10-A03) and approved by the Animal Ethics Committee of Guangzhou Laboratory. Mice were housed in a specific pathogen-free (SPF) environment under a 12-h light-dark cycle in the animal facility of Guangzhou National Laboratory, where the temperature was controlled at 18–23 °C and humidity was maintained at 40%–60%. Wild-type ICR mice were purchased from Charles River Laboratories (License number: SCXK (Yue) 2022-0063).

### Mouse embryonic nasal sample collection and dissociation

Timed-pregnant female ICR mice were euthanized by cervical dislocation at various embryonic days (E10.5 to E18.5) at 24-h intervals. The embryos were collected from the uterus, and nasal tissue was carefully dissected under a stereomicroscope and washed twice with cold PBS for single-cell suspension. The tissues were digested in 10 U/mL Papain (Worthington, LS003126) for 5 min at 37 °C, followed by mincing approximately 50 times to facilitate further dissociation. Subsequently, 100 U/mL DNase I (Worthington Biochemicals, LS006343) was added and the mixture was incubated for an additional 5–25 min. The enzymatic reaction was then terminated with 10% fetal bovine serum (FBS, NTC, #SFBE), followed by supplementation with 20 U/mL DNase I to ensure complete digestion of genomic DNA. After enzymatic digestion, cells were filtered through 40-μ m cell strainers (Falcon, 352360) on ice and centrifuged at 300 × *g* for 5 min at 4 °C. Cell counting and viability assessments were performed via trypan blue staining. The resulting cells were collected into PCR tubes for subsequent single-cell library preparation and sequencing.

### scRNA-seq library construction and sequencing

Single-cell suspensions from E10.5 to E18.5 embryos were loaded onto 10× Genomics Chromium Single Cell 3’ v3.1 system to generate single-cell gel beads-in-emulsion (GEMs). After incubation, the GEMs were disrupted to release the encapsulated single cells, and cDNA was subsequently synthesized and amplified via PCR. For library preparation, 25% of total cDNA (10 μL) was used to construct 3’ gene expression libraries. These libraries were then purified using SPRIselect beads (Beckman, B23319) to remove residual remaining reagents and contaminants, ensuring high-quality library preparation. Quality control of the libraries was performed using the Qsep100 system (Bioptic) for size distribution and Qubit 4.0 fluorometer for concentration quantification of the libraries. The sequencing of the libraries was carried out on DNBSEQ-T7 system (MGI), utilizing a paired-end mode (PE150) with a total output of 120 Gb of data.

### Preprocessing of scRNA-seq data

Raw FASTQ files from single-cell libraries generated using an MGI DNBSEQ-T7 system were processed as follows. Initial quality assessment was performed with fastp (v0.20.0). For sequence trimming, Trim Galore (v0.6.6) (integrated with Cutadapt) was utilized to perform hard-trimming, with the R1 and R2 reads were trimmed to 28 bp and 98 bp respectively (--hardtrim5). The processed clean reads were then aligned to the mm10 reference genome with mouse gene annotation (vM21) using STARsolo module within STAR (v2.7.6a) (Dobin et al., 2013). Alignment parameters were as follows: --soloType Droplet --soloFeatures Gene --soloCBstart 1 --soloCBlen 16 --soloUMIstart 17 -- soloUMIlen 12.

The Python library Scanpy (v1.6.0) (Wolf et al., 2018) was used to read and integrate the combined gene-barcode matrices of these samples. To ensure a high-quality cell atlas for downstream analysis, stringent quality control (QC) was performed based on the number of detected genes (nGenes), unique molecular identifiers (UMIs) and the percentage of mitochondrial genes (percent.mito). To account for both technical and biological variability across stages and replicates, sample-specific filtering thresholds were applied (see Table S1 for details). Generally, low-quality cells with high mitochondrial content were excluded. Furthermore, potential doublets were identified and removed using Scrublet (v1.0) (Wolock et al., 2019) with scanpy.pp.scrublet. Following these filtering steps, a total of 183,247 high-quality cells were retained, with a median of 3,777 detected genes and 12,822 UMIs per cell.

### Cell cluster identification

To minimize technical variability, major cell clusters were identified for each sample separately prior to dataset integration. Raw counts were normalized to 10,000 counts per cell and log-transformed. The top 3,000 highly variable genes (HVGs) were selected for downstream analysis. Principal component analysis (PCA) was performed on these HVGs, and the first 30 principal components were used to construct a k-nearest-neighbor (KNN) graph (n_neighbors = 10). Uniform Manifold Approximation and Projection (UMAP) was employed for visualization. Cell clusters were identified using the Leiden graph-clustering algorithm. To characterize cluster identities, differentially expressed genes (DEGs) were identified using the Wilcoxon rank-sum test. Significant marker genes were defined by a Benjamini–Hochberg adjusted *P*-value < 0.01 and a log fold change (log_2_FC) ≥ 0.25.

To achieve a high-resolution and biologically accurate characterization of the cellular landscape, a hierarchical annotation strategy was employed based on canonical markers curated from published literature (Table S2). Initial broad clusters were annotated by cross-referencing the expression of these markers, and their specificity was verified using dot plots and UMAP feature plots. An iterative sub-clustering strategy was then performed to refine annotations. First, clusters expressing proliferation-related genes (e.g., *Top2a*, *Mki67*) were labelled with the suffix “_prolif” (e.g., “Chondrocyte_prolif”). Second, heterogeneous populations were further stratified through re-clustering. For example, ciliated cells were subdivided into “Ciliated cells1_Sftpc−” and “Ciliated cells2_Sftpc+” subpopulations based on differential expression of *Sftpc* and *Cdhr3*. Third, low-abundance populations, including *Gnrh1*^+^ neurons, *Prokr2*^+^ terminal neurons, and *Sox10*^+^/*Foxd3*^+^ neural crest cells, were identified after dataset integration to improve detection sensitivity. This process resulted in 52 distinct cellular clusters.

Additionally, gene ontology (GO) enrichment analysis for DEGs associated with each cluster was performed using the clusterProfiler R package (Yu et al., 2012) in R.

### Data Integration

To characterize the global cellular landscape and the epithelial compartment, dimensionality reduction was performed based on feature selection. HVGs were identified with expression thresholds (min_mean = 0.0125, max_mean = 3, and min_disp = 0.5) to capture biological variability. PCA was conducted on these HVGs, and the top 40 PCs for global atlas or 30 PCs for epithelial subset were retained. The neighborhood graph was then constructed (n_neighbors = 10), followed by UMAP visualization.

For the mesenchymal lineage, Harmony (Korsunsky et al., 2019) was used to correct for batch effects. The top 2,000 HVGs were selected with (batch_key = “Batch”). After computing the top 30 PCs, batch-corrected embeddings were generated by Harmony and used for neighborhood graph construction and UMAP visualization.

For several regions of neurogenic and non-neurogenic epithelial cells, batch correction was performed using BBKNN (Batch-Balanced k-Nearest Neighbors) algorithm (Polanski et al., 2020). The top 2,000 HVGs were identified using batch-aware selection, followed by PCA (30 components) and BBKNN correction. To visualize continuous developmental trajectories, PHATE (Potential of Heat-diffusion for Affinity-based Trajectory Embedding) (Moon et al., 2019) was applied. PHATE embeddings of respiratory epithelial cells were generated with parameters (n_components = 2, knn = 40, gamma = 0.3), optimized to preserve global trajectory structures.

### Pseudo-bulk RNA analysis

Pseudo-bulk RNA expression profiles were generated by aggregating single-cell raw counts corresponding to each time point. Counts per million (CPM) normalization followed by log transformation was performed. PCA was performed on the log-CPM matrix using prcomp function, and the first two principal components were extracted for visualization. To model temporal gene expression dynamics, a smooth curve was fitted to the aggregated profiles using the principal_curve function from the R package princurve (v2.1.6). Subsequently, the PCA scatter plots with overlaid fitted curves were visualized using ggplot2 (v3.4.0).

### RNA velocity analysis

To infer directional state transitions, spliced and unspliced RNA transcripts were quantified for each sample using the STARsolo algorithm with the Velocyto option enabled. RNA velocity was estimated for mesenchymal and epithelial populations using scVelo (v0.2.3) (Bergen et al., 2020). Specifically, normalized moments of spliced and unspliced counts were calculated using 30 principal components and 20 nearest neighbors. The dynamical model was then applied to recover transcriptional kinetics, and RNA velocity was subsequently estimated in dynamical mode. A velocity graph was then constructed using 20 nearest neighbors, and velocity vectors were projected onto the corresponding low-dimensional embeddings for visualization.

### Trajectory inference

For mesenchymal trajectory inference, Partition-based Graph Abstraction (PAGA) (Wolf et al., 2019) was applied to reconstruct transitions between clusters. In this framework, edge weights represent connectivity strengths inferred from velocity-based transitions. Developmental progression was further evaluated by diffusion pseudotime (DPT) analysis.

For the respiratory epithelium, PAGA analysis was performed following the same strategy used for mesenchymal cells. In addition, Monocle2 (Qiu et al., 2017) was used to independently reconstruct the trajectory of respiratory epithelium cells. Specifically, the top 2,000 HVGs were selected to capture relevant transcriptional variability. Dimensionality reduction was conducted using the DDRTree algorithm, followed by cell ordering to define lineage relationships.

To further investigate the temporal dynamics of GBC subtypes, the dataset in this study was integrated with three published nasal epithelium datasets from GEO: P19 from GSE224604 (Yang et al., 2024), P8W from GSE185168 (Horgue et al., 2022), and P11W-P36W from GSE245074 (Ualiyeva et al., 2024). After integration, Monocle2 was applied to construct trajectories of GBC subtypes using 2,000 HVGs.

### TF activity inference

Regulon inference and activity analysis were performed using the pySCENIC framework (v0.11.0) (Aibar et al., 2017) to identify transcription factor (TF)-centered regulatory programs across defined cell subpopulations. The analysis was conducted on a filtered expression matrix comprising 6,000 HVGs. First, co-expression relationships between TFs and potential target genes were inferred using the GRNBoost2 algorithm with mouse TFs. The resulting TF-target adjacencies were grouped into co-expression modules. To obtain high-confidence regulons, cis-regulatory motif enrichment analysis was performed using prune2df based on the mm10 motif databases. Only TF-target interactions supported by significant motif enrichment (normalized Enrichment Score, NES > 3.0) were retained.

Regulatory activity at the single-cell level was quantified using AUCell. To improve the robustness, several filtering steps were applied: (i) TFs with low expression or activity (mean expression < 0.1) were excluded; (ii) regulons showing no activity across cell groups were removed; and (iii) regulons with motif enrichment NES > 3 were retained for downstream analysis. The final mean AUCell activity matrix was visualized using the ComplexHeatmap (v2.16.0) R package, with hierarchical clustering performed using the ward.D2 method to reveal cell-type-specific regulatory patterns.

### Cell-cell interaction analysis

To characterize the intercellular communication landscape during mouse nasal development, the CellChat framework (v1.6.1) (Jin et al., 2021) was applied. Interactions were inferred based on the CellChatDB.mouse ligand-receptor database. Overexpressed ligands and receptors in each cell group were first identified, after which communication probabilities between cell groups were computed using a permutation-based approach implemented in CellChat. To ensure statistical robustness, only cell groups containing at least 10 cells were included in the analysis. Pathway-level communication probabilities were subsequently calculated by aggregating individual ligand–receptor pairs. To focus on high-confidence interactions and minimize redundancy, only ligand–receptor pairs with a communication probability greater than 0.01 were retained. When multiple interactions were detected between the same sender–receiver pair, only the interaction with the highest probability was kept for visualization. The overall signaling strengths and specific ligand-receptor interactions across subpopulations were visualized using bubble plots and circle plots.

### Bulk RNA library preparation and sequencing

Total RNA was extracted using TRIzol Reagent (MRC, TR118) following the manufacturer’s instructions. Subsequently, sequencing libraries were constructed using the VAHTS mRNA-seq v2 Library Prep Kit for Illumina (Vazyme, NR605). Library sequencing was performed on an Illumina NovaSeq X Plus platform (Annoroad Gene Technology).

### Bulk RNA-seq analysis

The paired-end sequenced reads were aligned to a transcriptome index generated from the GENCODE transcriptome annotation (M13) using Bowtie2, and transcript quantification was performed with RSEM (Li and Dewey, 2011). The resulting gene expression profiles were output in the *gene.results* file. Raw read counts were used for differential expression analysis, while TPM values were used for expression visualization and comparison. Differentially expressed genes (DEGs) were identified using edgeR within the R/Bioconductor framework (v3.6.1), applying a false discovery rate (FDR) < 0.05 and an absolute log₂ fold change cutoff. GO enrichment analysis was performed with DEGs using the R package clusterProfiler, with FDR correction applied for multiple testing. GO terms with an adjusted *P*-value < 0.05 were considered significantly enriched. Figures were generated using the ggplot2 package (v3.3.2).

### Primary RE progenitor cells cultures

Timed-pregnant ICR mice at E11.5 were euthanized by cervical dislocation. Embryos were collected and washed twice with cold PBS. Nasal tissues were carefully dissected from embryos under a stereomicroscope using fine forceps. Dissected tissues were incubated in 20U/mL Dispase II (Gibco, #17105-041) for 10 min to facilitate separation of the epithelial layer from the underlying mesenchyme. The respiratory epithelial (RE) progenitor region was identified based on FGF8 immunostaining in E11.5 nasal tissue sections, as previously described (Kawauchi et al., 2005). The *Fgf8*^+^ RE progenitor region was then microdissected using insulin syringe needles (BD, #328421). The isolated epithelium was subsequently fragmented into small clusters by treatment with 10 U/mL Papain (Worthington, LS003126) at 37 °C for 1 min, and terminated by adding an equal volume of 10% FBS. Cell clusters were embedded in Matrigel (Corning, #354234) and cultured in ALL medium. Cultures were maintained at 37 °C in a humidified with 5% CO2 atmosphere, with medium refreshed every 48 h in 24-well plate.

### Foxa1 overexpression

The full-length coding sequence (CDS) of mouse *Foxa1* was amplified by PCR from cDNA derived from mouse lung tissue, and cloned into the pMXs expression vector. Retroviruses were produced in Plat-E cells via polyethylenimine (PEI)-mediated transfection. Briefly, 7.5 × 10^6^ Plat-E Cells were seeded in 10**-**cm dish and cultured in medium containing 10% FBS for 12-18 h before PEI transfection. For transfection, 20 μg plasmid DNA was mixed with 40 μL PEI reagent (#MW40000, YEASEN) in Opti-MEM and incubated for 15 min at room temperature. The mixture was then added to the Plat-E cells, and the medium was replaced 10 h post-transfection. Retroviral supernatants were harvested 48 h post-transfection.

For 3D culture, 80 μL of ice-cold Matrigel was first added to each well of a 48-well plate, evenly covering the well bottom, and allowed to solidify at 37 °C for 20 min. Subsequently, 400 μ L of the cell–virus mixture was added to the Matrigel-coated wells and incubated for 16 h at 37 °C. Non-viable cells were then removed by aspiration, followed by addition of a 60 μL Matrigel overlay and solidification at 37°C for 20 min. Cultures were then maintained in ALL medium, with medium changes every 48 h.

### Single-molecule fluorescence in situ hybridization (smFISH)

SmFISH was performed following the protocol as previously described (Cao et al., 2023). Tissue sections (10 μm) were cut using a Leica CM3050S cryostat at –20 °C, and mounted onto adhesive glass slides (CITOTEST, #80312-3161). Sections were fixed in 4% paraformaldehyde (PFA) in PBS for 10 min at room temperature, washed three times with PBS (5 min each), and then immersed in 70% (vol./vol.) ethanol for at least 1 h at room temperature.

The unlabeled gene-specific primary probes were synthesized by Sangon Biotech (China). Secondary FLAP probes, conjugated with Cy3 or Cy5 fluorophores via 5’-amino modifications, were purchased from Thermo Fisher Scientific (China). The sequences of all probes are provided in Table S9.

For in vitro pre-annealing, a reaction mixture containing 8.5 μL primary probe mix, 0.5 μL secondary FLAP probe (Cy3 or Cy5), and 1 μL 10×NEB 3 buffer was prepared on ice, followed by gentle pipetting to mix. The mixture was subjected to thermal cycling with the following program: 85°C for 3 min; cooling to 65 °C at –1 °C/s and holding for 3 min; further cooling to 25 °C at –0.1 °C/s and incubating for 5 min. After cycling, the mixture was gently mixed by pipetting without vortexing.

Hybridization buffer containing 10% formamide (DFA) was prepared in a fume hood. The pre-annealed probe duplexes were diluted 1:50 (v/v) in the hybridization buffer to a final working concentration of 3.2 nM for subsequent in situ hybridization. For combined smFISH and immunofluorescence staining, mouse monoclonal anti-E-cadherin (CDH1, 1:200; BD Transduction Laboratories, 610181) was added into diluted probe-hybridization buffer. The tissue sections on slides were outlined with a PAP pen, and Wash Buffer A (WA, Biosearch Technologies, #SMF-WA1-60) was added dropwise to the circled area for rehydration at room temperature for 10 min. The WA was then carefully removed, and the diluted probe-hybridization buffer with primary antibody was added dropwise onto each section individually. The slides were placed in a humidity-controlled chamber and incubated at 37 °C for 16 h in the dark. After hybridization, slides were washed twice by pre-warmed Wash Buffer A with Alexa Fluor-conjugated donkey anti-mouse secondary antibody (488, 1:400, Thermo Fisher) (37°C, 30 min each), then stained with 200 ng/mL DAPI in Wash Buffer B (Biosearch Technologies, #SMF-WB1-20) for 5 min, and given a final wash with Wash Buffer B. Sections were mounted with 50-100 μL anti-fade medium under coverslips, with edges sealed using clear nail polish.

Confocal images were acquired using Olympus IXplore SpinSR confocal microscope. Image processing including brightness/contrast adjustment, pseudo-coloring, and z-stack alignment was performed uniformly across datasets using Adobe Photoshop CC 2020 and Fiji/ImageJ software.

### Immunofluorescence

Heads from ICR mouse embryos at embryonic days 11.5, 14.0 and 18.0 (E11.5, E14.0, and E18.0) were dissected and fixed in 4% paraformaldehyde (PFA) for 16 h. Following fixation, the samples were dehydrated overnight in 30% sucrose at 4 °C. The head tissue was embedded in OCT compound (Sakura, #4583) and cryosectioned into 10 μm-thick sections. Sections were mounted onto adhesive glass slides and stored at 4 °C until further processing.

For immunofluorescence staining, sections were washed three times with PBS, permeabilized with 0.5% Triton X-100 for 20 min, and then blocked at room temperature for 1 h with blocking buffer (5% BSA in PBS). Following blocking, section were incubated overnight at 4 ° C with primary antibodies: rabbit polyclonal anti-SFTPC (1:200, Invitrogen, PA5-102493) and mouse monoclonal anti-acetylated-α-Tubulin(1:1000, Sigma, T7451). Subsequently, sections were incubated with Alexa Fluor-conjugated donkey or goat anti-mouse or anti-rabbit secondary antibodies (488/568, 1:400, Thermo Fisher) for 2 h at room temperature. Nuclei were visualized with DAPI (Invitrogen, D1306) for nuclear staining, and slides were mounted using appropriate anti-fade mounting media (KeyGEN BioTECH, KGA1523-5). Images were captured using the Andor Dragonfly 200 confocal microscope.

### Quantitative real-time PCR (qRT–PCR)

Total RNA was isolated using TRIzol Reagent (MRC, TR118). Complementary DNA (cDNA) was synthesized from 1 μg of RNA via reverse transcription with dNTPs (TIANGEN, CD111-03), oligo (dT) 18 (TakaRa, 3806), RT Ace (TOYOBO, TRT-101), RNase inhibitor (Promega, N2511) and 5×RT Buffer (TOYOBO, TRT-101). The qRT–PCR was performed using an iTaq™ Universal SYBR® Green Supermix (Bio-Rad #1725124) on a CFX Connect Real-Time PCR System (Bio-Rad). *Gapdh* was used as the endogenous control. Relative gene expression was calculated using the ΔΔCt method. All reactions were performed in technical triplicate, with no-template controls included. For statistical analysis, two-tailed unpaired Student’s *t*-test were used for comparisons between two groups, while ordinary one-way ANOVA followed by Dunnett’s post hoc test was used when involving multiple groups (GraphPad Prism 8). Significance was defined as *P* < 0.05. All primer sequences are provided in Table S10.

### Data availability

"The datasets generated in this study have been deposited in GEO and will be made publicly available upon publication.

## CRediT authorship contribution statement

**Huan Chen**: Methodology, Data analysis, Organoid culture, Manuscript writing. **Yingxiu Chen**: Methodology, Library generation, smFISH, Manuscript writing. **Mengjie Pan**: Methodology, Embryonic nose dissection. **Ziyu Feng**: Methodology, Nose tissue dissociation. **Baomei Cai**: Methodology, Library generation. **Yiyi Cheng**: Organoid culture. **Sihao Chen, Jiehong Deng, Xia Yao, Chunhua Zhou, Yunjing Du, Wei He, Ruifang Zhang, Yudong Fu, Shujuan Liu**: Technical and experimental support. **Lihui Lin, Shengyong Yu, Yuehong Yan, Duanqing Pei, Dajiang Qin, Jiekai Chen**: Advices. **Dajiang Qin, Jiekai Chen**: Funding acquisition. **Shangtao Cao**: Conceptualization, Manuscript writing, Funding acquisition, Supervision

## Conflict of interest

The authors declare that they have no conflict of interest.

## Acknowledgement

This research was supported by grants from the National Natural Science Foundation of China (82470704); Major Project of Guangzhou National Laboratory (GZNL2023A02005); the National Key R&D Program of China (2024YFA1802300); Science and Technology Planning Project of Guangdong Province, China (2023TQ07A630); Key Laboratory of Guangdong Higher Education Institutes (2021KSYS009); Guangzhou Key Laboratory of Biological Targeting Diagnosis and Therapy (202201020379); We also thank a lot for public instrument service center Guangzhou National Laboratory and the support from the Guangzhou Branch of the Supercomputing Center of the Chinese Academy of Sciences.

## Notes

### Competing Interest Statement

The authors have declared no competing interest.

### Summary of Updates

The manuscript has been revised to update the Data Availability statement. The associated datasets are undergoing further processing and will be made publicly available upon publication. Minor textual revisions have also been made for clarity.

