## supplement figure for "A Single-Cell Temporal Atlas of Mouse Nasal Embryonic Development"

### Figure S1 related to Fig.1

A

| | E10.5 | E11.5 | E12.5 | E13.5 | E14.5 | E15.5 | E16.5 | E17.5 | E18.5 |
|---|---|---|---|---|---|---|---|---|---|
| Batch | 2 | 2 | 2 | 2 | 2 | 2 | 2 | 2 | 2 |
| Embryo | 4 | 2 | 2 | 2 | 2 | 2 | 2 | 2 | 2 |
| nCells | 17,966 | 20,467 | 18,223 | 15,785 | 25,478 | 20,927 | 20,927 | 22,906 | 19,666 |

Image

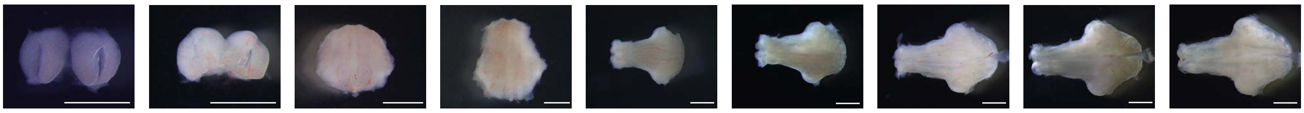

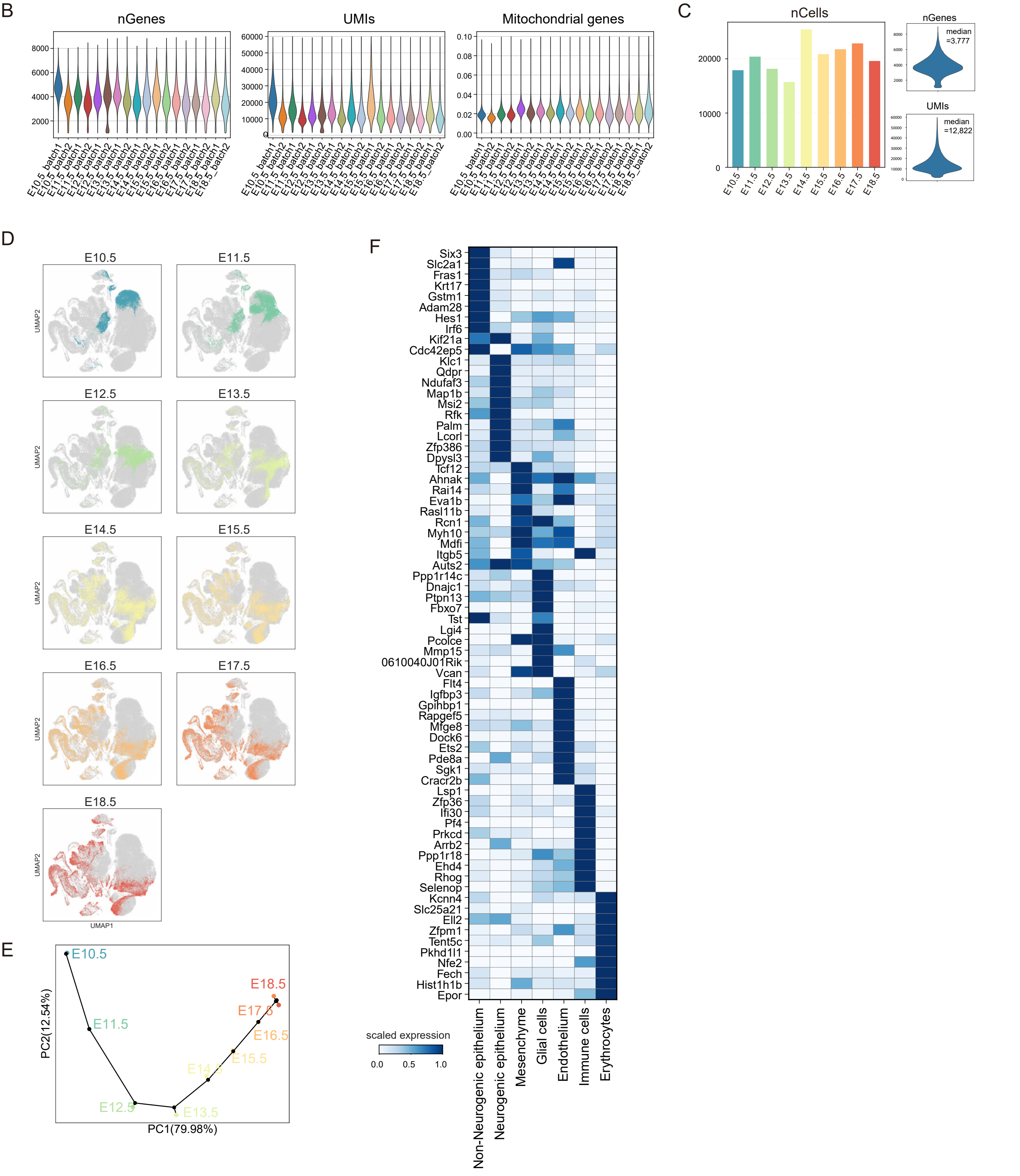

**Fig. S1. Tissue morphology and data quality of mNEDCA (related to Fig. 1)**

A: Table summarizing the number of experimental replicates (batch), embryos, and total high-quality cells (nCells) collected at each stage. Representative images of dissected mouse nasal tissue across developmental stages from E10.5 to E18.5 are shown in the bottom. Scale bars,1 mm.

B: Quality control metrics for single-cell RNA sequencing data. Violin plots representing the number of detected genes (Genes), unique molecular identifiers (UMIs) and the percentage of mitochondrial genes from each sample. Quality control and filtering metrics are detailed in Table S1.

C: Left, histograms showing the number of high-quality cells per embryonic stage. Right, violin plots summarizing the number of Genes and UMIs for all cells. Median numbers of Genes and UMIs are indicated above the plots.

D: UMAP visualization showing the distribution of cells colored by each embryonic period. In each panel, cells from the indicated stage are highlighted in color, while other cells are shown in gray.

E: Principal component analysis (PCA) of pseudo-bulk RNA profiles aggregated by developmental time based on scRNA-seq data. PC1 corresponds to developmental progression, indicating continuity and concordance among samples.

F: Matrix plot showing normalized expression of the differentially expressed genes across major cell types. Color scale indicates row-normalized expression levels.

Figure S2 related to Fig.1

A

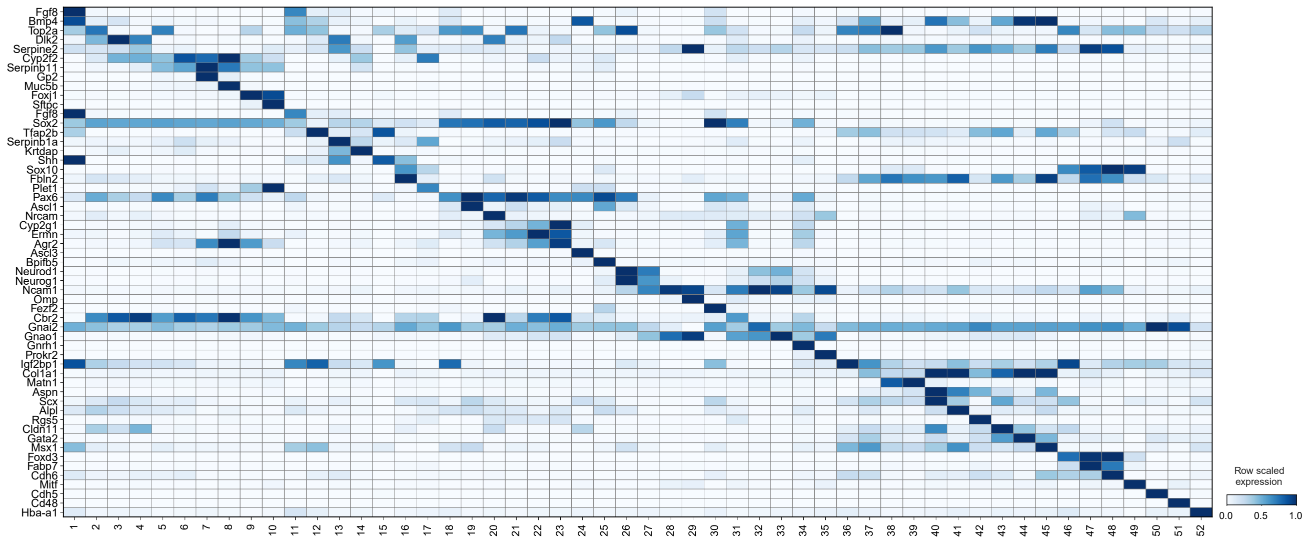


B

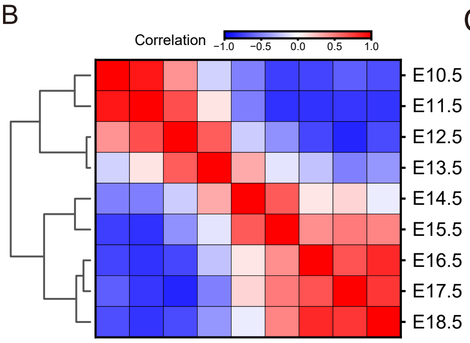


C

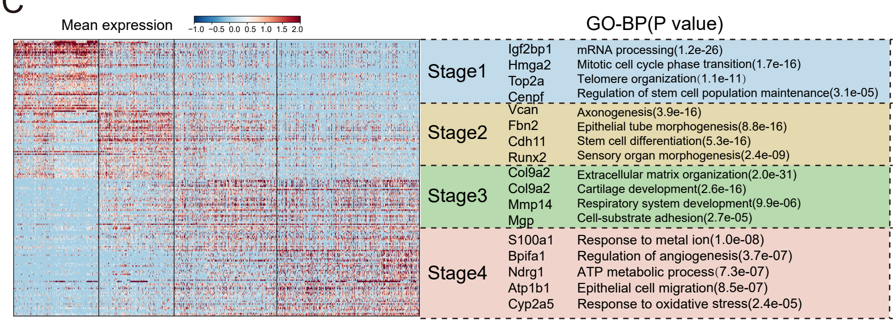

**Fig. S2. Single-cell transcriptomic landscape and stage-resolved molecular dynamics of mNEDCA (related to Fig. 1)**

A: Matrix plot showing normalized expression of the representative marker genes across 52 cell types. Color scale shows row-normalized expression levels.

B: Heatmap showing pairwise Pearson correlations between embryonic period based on scRNA-seq expression profiles. Hierarchical clustering (left dendrogram) partitions the developmental timeline into four major groups.

C: Left, heatmap of differentially expressed genes (DEGs) across the four presumptive developmental stages (Wilcoxon rank-sum test, *P* value < 0.01), selected genes are labelled. Genes are row-scaled to emphasize relative expression differences across stages. Right, selected top Gene Ontology (GO) biological process terms enriched among stage-specific DEGs, with enrichment significance indicated by *P*-values.

Figure S3 related to Fig.2

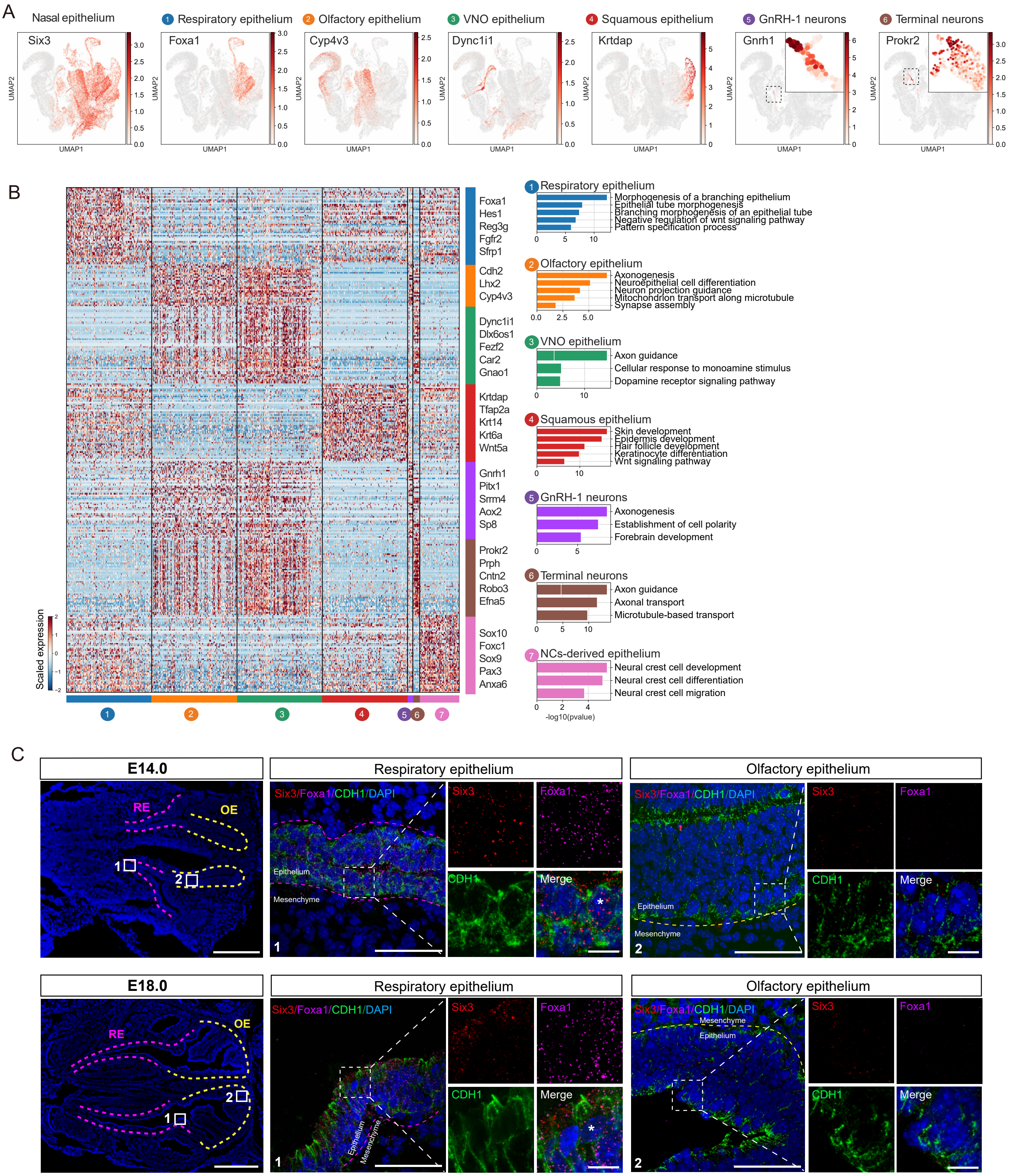

**Fig. S3. Region-specific gene expression profiles in the neurogenic and non-neurogenic epithelium (related to Fig. 2)**

A: UMAP projection showing expression of differentially expressed markers across several regions. *Six3* marks nasal epithelium, *Foxa1* marks respiratory epithelium, *Cyp4v3* marks olfactory epithelium, *Dync1i1* marks VNO epithelium, *Krtdap* marks squamous epithelium, *Gnrh1* marks GnRH-1 neurons and *Prokr2* marks terminal neurons.

B: Left, heatmap of differentially expressed genes among the several regions of the epithelial populations (Wilcoxon rank-sum test, $P$ value < 0.01), selected genes are labelled. Right, bar plot displaying GO terms enriched among these genes, with enrichment significance indicated by $P$-values.

C: SmFISH combined with immunofluorescence staining of nasal tissue at E14.0 (top) and E18.0 (bottom). Low-magnification scans (left) provide an overview of respiratory and olfactory regions. High-magnification views showing specific *Foxa1* (magenta) expression restricted to respiratory epithelium (middle; asterisks denote individual positive cells), and absent from olfactory epithelium (right). *Six3* (red) marks nasal epithelium and CDH1 (green) immunofluorescence outlines epithelial boundaries. Images are representative of at least three independent experiments. Scale bars: 500 μm (left), 40 μm (middle/right); 10 μm for insets.

### Figure S4 related to Fig.2

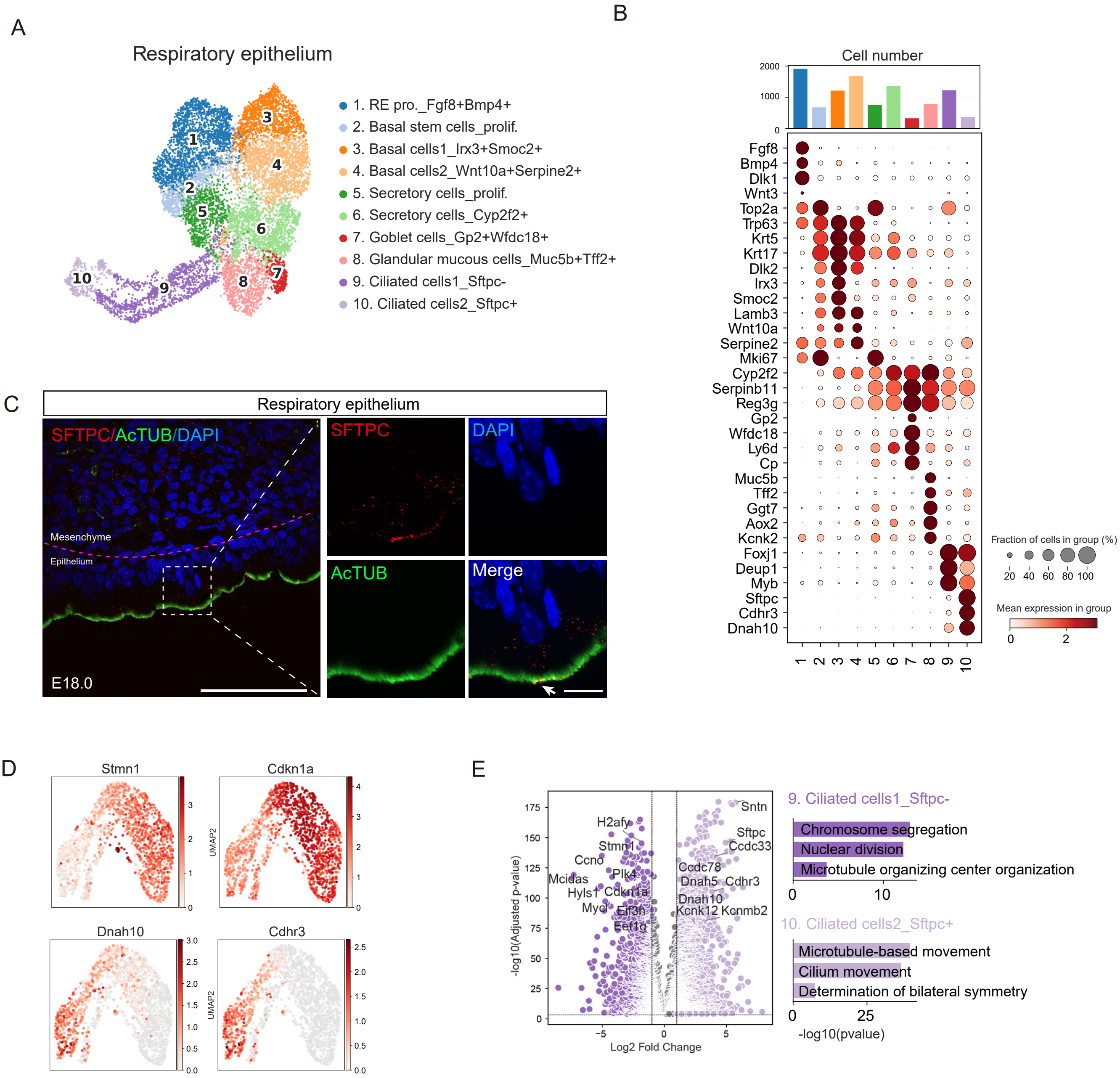

**Fig. S4. Heterogeneity of respiratory epithelial cells and two ciliated subtypes (related to Fig. 2)**

A: UMAP visualization of respiratory epithelial cells categorized by cell types (BBKNN integration, n_pcs = 30, n_neighbors = 10). Each dot represents one cell.

B: Dot plot representing the expression of marker genes associated with different cell types in the respiratory epithelium. Dot color indicates average normalized expression and dot size indicates the fraction of cells expressing each gene. The bar plot above indicates cell counts per cluster.

C: Immunofluorescence staining of SFTPC (red) and acetylated α-tubulin (AcTUB, green) within respiratory epithelium of E18.0 nasal tissue. Arrowhead indicates ciliated cells2_Sftpc+ subtype. Images are representative of at least three independent experiments. Scale bars: 60 μm; 10 μm for insets.

D: UMAP feature plots showing expression of ciliated cells1_Sftpc$^{-}$ marker (*Stmn1*, *Cdkn1a*) and ciliated cells2_Sftpc$^{+}$ marker (*Dnah10*, *Cdhr3*).

E: Left, volcano plot showing genes differentially expressed between the two ciliated subclusters (Wilcoxon rank-sum test, *P* value < 0.01). Right, bar plot displays GO terms enriched among these genes, with enrichment significance indicated by *P*-values.

### Figure S5 related to Fig.2

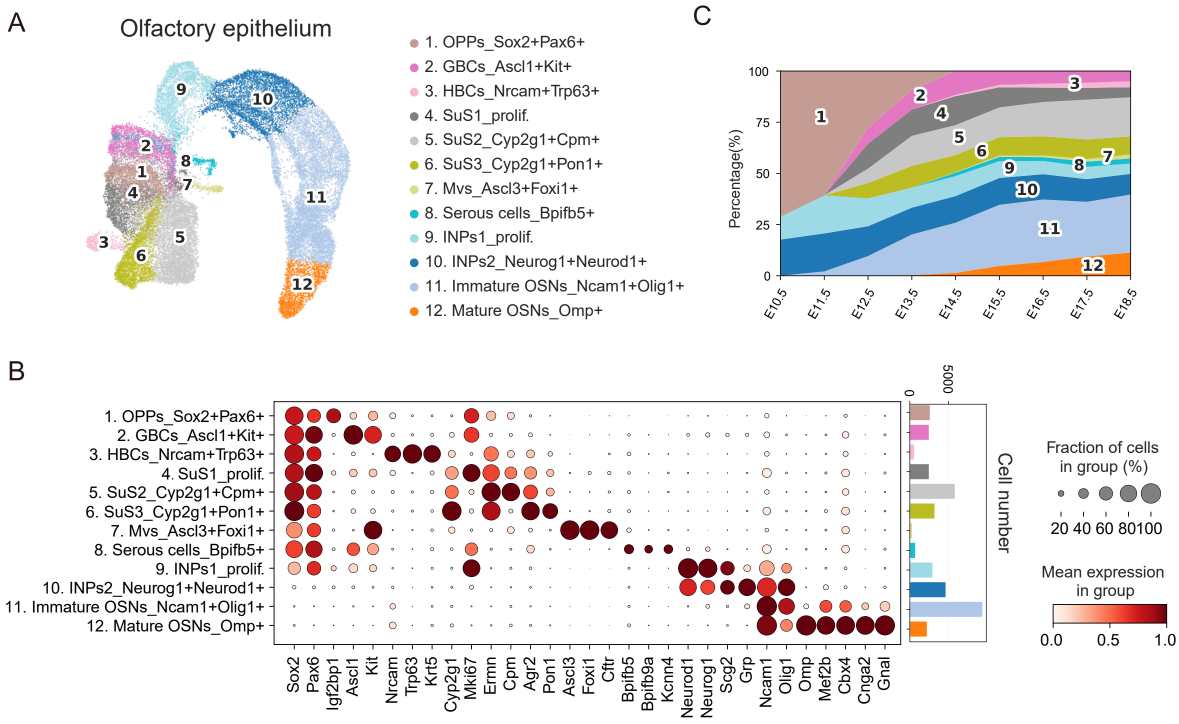

**Fig. S5. Heterogeneity of olfactory epithelial cells (related to Fig. 2)**
A: UMAP visualization of olfactory epithelium (37,584 cells) categorized by cell type (BBKNN integration, n_pcs = 30, n_neighbors = 10). Each dot represents one cell.
B: Dot plot showing expression of marker genes associated with different cell types in the olfactory epithelium. Dot color indicates average normalized expression and dot size indicates the fraction of cells expressing each gene. The bar plot right represents the cell counts per cluster.
C: Stacked plot showing the fraction of each cell type across each embryonic stage. Colors correspond to panel (A).

### Figure S6 related to Fig.2

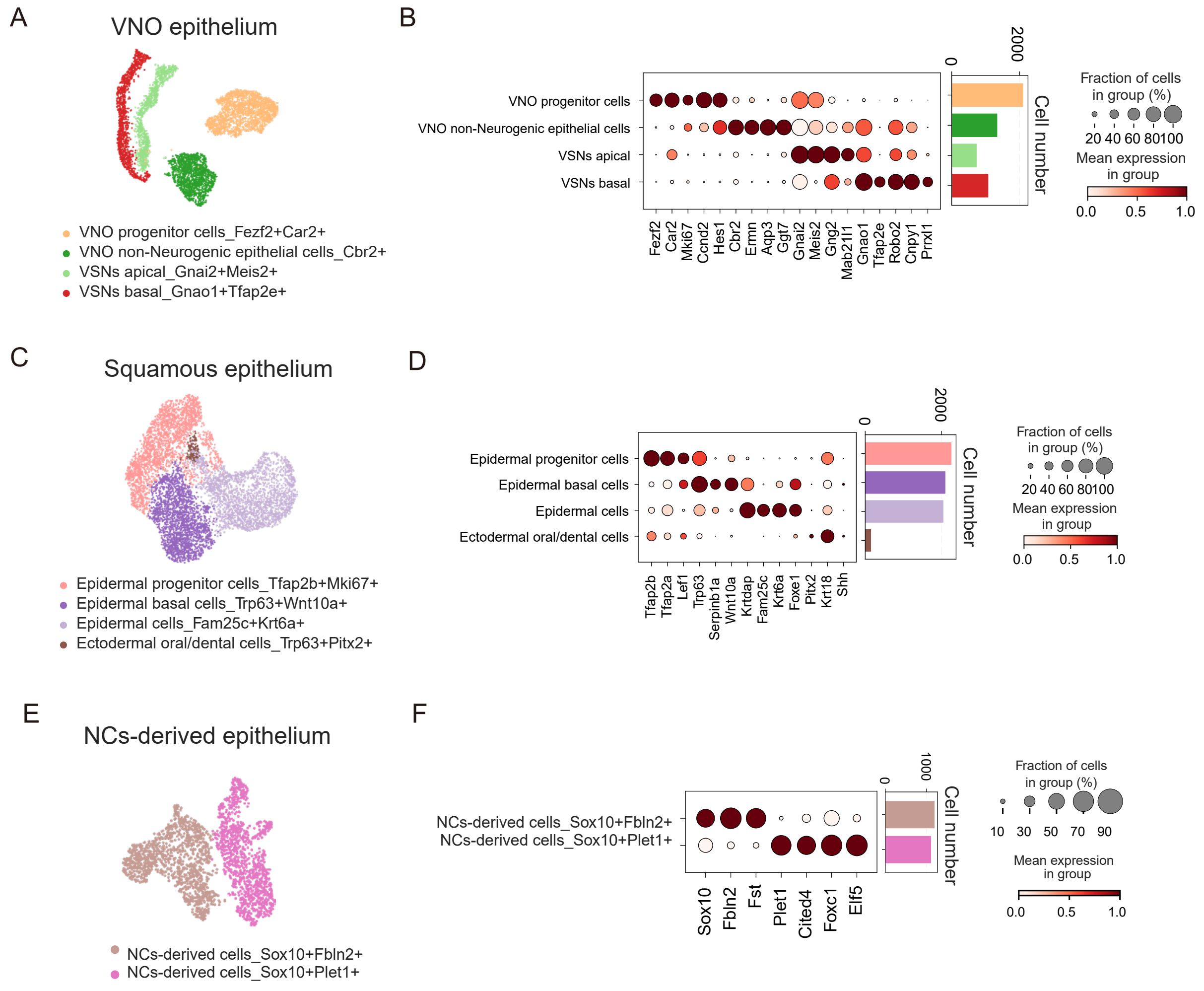

**Fig. S6. Annotation of cell types in different regions of epithelial populations (related to Fig. 2)**

A: UMAP visualization of VNO epithelium (5,877 cells) categorized by cell type (BBKNN integration, n_pcs = 30, n_neighbors = 10). Each dot represents one cell.

B: Dot plot showing expression of marker genes associated with cell types in VNO epithelium. Dot color indicates average normalized expression and dot size indicates the fraction of cells expressing each gene. The bar plot right represents the cell counts per cluster, and colors correspond to panel (A).

C: UMAP visualization of squamous epithelium (6,608 cells) categorized by cell type (BBKNN, n_pcs = 30, n_neighbors = 10). Each dot represents one cell.

D: Dot plot showing expression of marker genes associated with cell types in squamous epithelium. Dot color indicates average normalized expression and dot size indicates the fraction of cells expressing each gene. The bar plot right represents the cell counts per cluster, and colors correspond to panel (C).

E: UMAP visualization of neural crests-derived cells (2,317 cells) categorized by cell type (BBKNN integration, n_pcs = 30, n_neighbors = 10). Each dot represents one cell.

F: Dot plot showing expression of marker genes associated with cell types in neural crests-derived cells. Dot color indicates average normalized expression and dot size indicates the fraction of cells expressing each gene. The bar plot right represents the cell counts per cluster, and colors correspond to panel (E).

### Figure S7 related to Fig.3

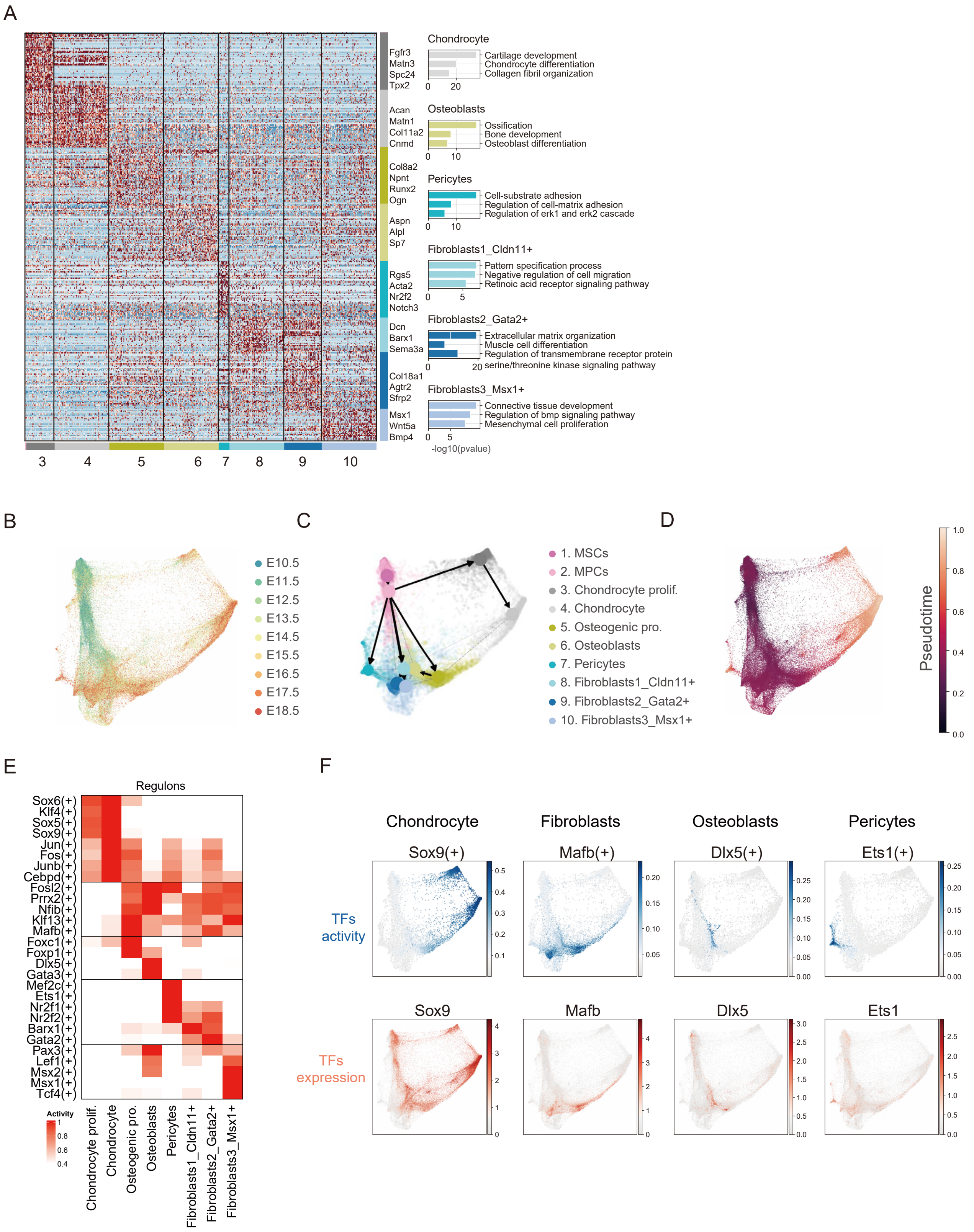

**Fig. S7. Functional characteristics and developmental trajectory of nasal mesenchyme (related to Fig. 3)**

A: Left, heatmap showing differentially expressed genes among cell types of nasal mesenchyme (Wilcoxon rank-sum test, $P$ value < 0.01), selected genes are labelled. Right, bar plot displaying GO terms enriched among these genes, with enrichment significance indicated by $P$-values.

B: PHATE visualization of nasal mesenchyme colored by embryonic stage. Each dot represents one cell. PHATE, Potential of Heat-diffusion for Affinity-based Trajectory Embedding.

C: PAGA graph abstraction overlaid on a PHATE map of the nasal mesenchyme. Node positions and connectivity were computed using Scanpy's PAGA (see Methods).

D: PHATE graph of nasal mesenchyme colored by Diffusion Pseudotime (DPT). The gradient from dark purple to bright yellow represents the predicted differentiation progression.

E: Heatmap of regulon activity (AUC scores) among differentiated cell types of nasal mesenchyme inferred by pySCENIC. AUC scores were calculated at the single-cell level and averaged per cell type. Regulons are row-z-score normalized. Regulons were filtered for measurable activity (see Methods).

F: UMAP showing the TF activity (blue, AUC score) and gene expression (red, log-normalized) of chondrocyte-specific (*Sox9*), fibroblasts-specific (*Mafb)*, osteoblasts-specific (*Dlx5)* and pericytes-specific (*Ets1)*. SCENIC-generated TF activity, represented by AUC score, reflects the co-expression strength of TF and its target genes.

Figure S8 related to Fig.4

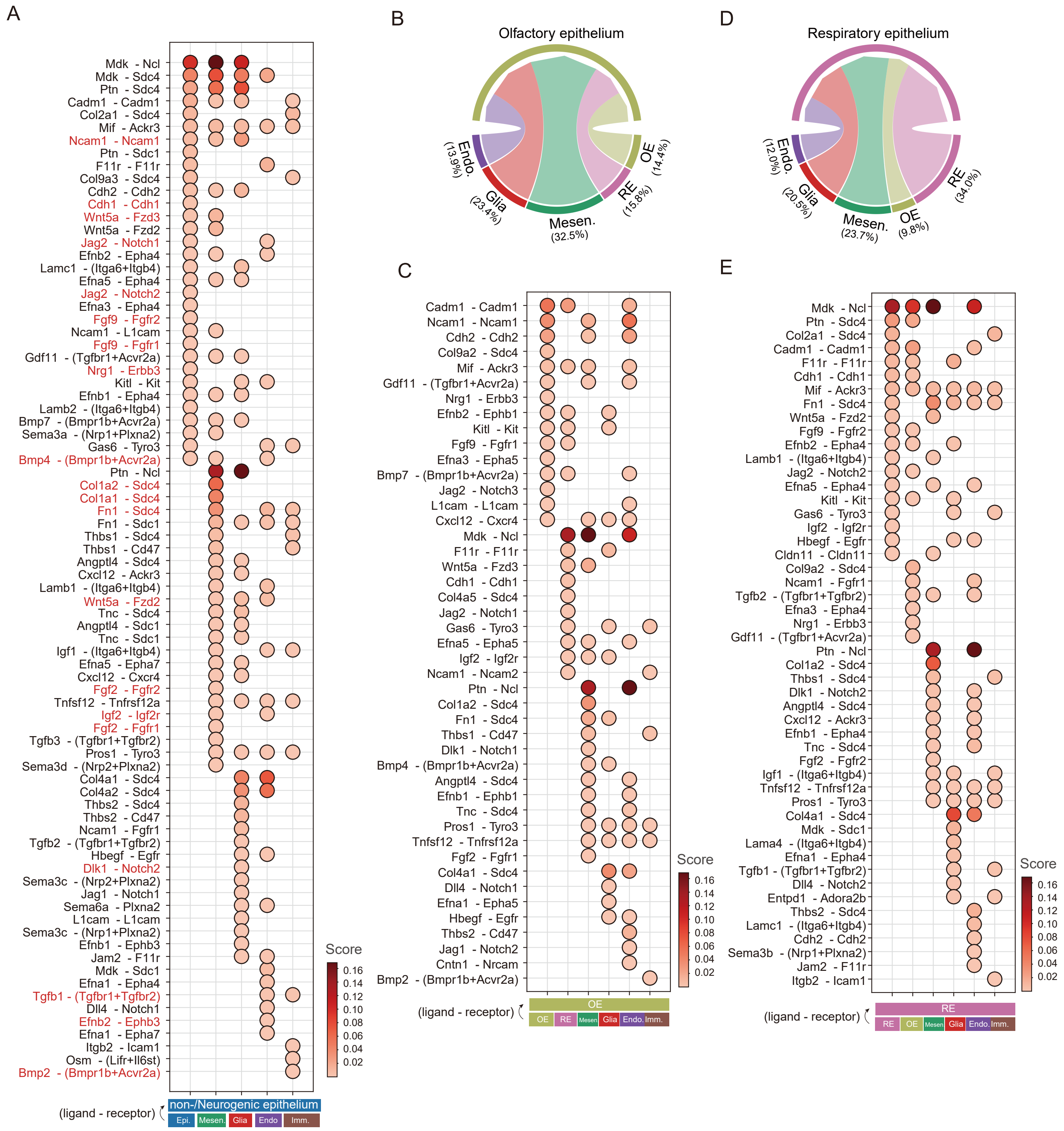

**Fig. S8. Cell-Cell interactions and signaling of nasal respiratory and olfactory epithelium development (related to Fig. 4)**

A: Dot plot showing LR interactions strength related to Fig. 4A. The color of each dot represents the communication probability. Only significant LR pairs are displayed (filtering criteria and analysis are detailed in Methods). Known signaling pathways are marked in red. Epi., non-/neurogenic epithelium; Mesen., nasal mesenchyme; Glia, glia cells; Endo., endothelium; Imm., immune cells.

B: Circos plot of LR interactions between OE and other cell types. Ribbon widths represent the interaction weight. OE, olfactory epithelium; RE, respiratory epithelium; Mesen., nasal mesenchyme; Glia, glia cells; Endo., endothelium; Imm., immune cells.

C: Dot plot showing LR interactions strength related to panel (B). The color of each dot represents the communication probability. Only significant LR pairs are displayed.

D: Circos plot of LR interactions between RE and other cell types. Ribbon widths represent the relative weight of LR interactions between compartments. OE, olfactory epithelium; RE, respiratory epithelium; Mesen., nasal mesenchyme; Glia, glia cells; Endo., endothelium; Imm., immune cells.

E: Dot plot showing LR interactions strength related to panel (D). The color of each dot represents the communication probability. Only significant LR pairs are displayed.

Figure S9 related to Fig.5

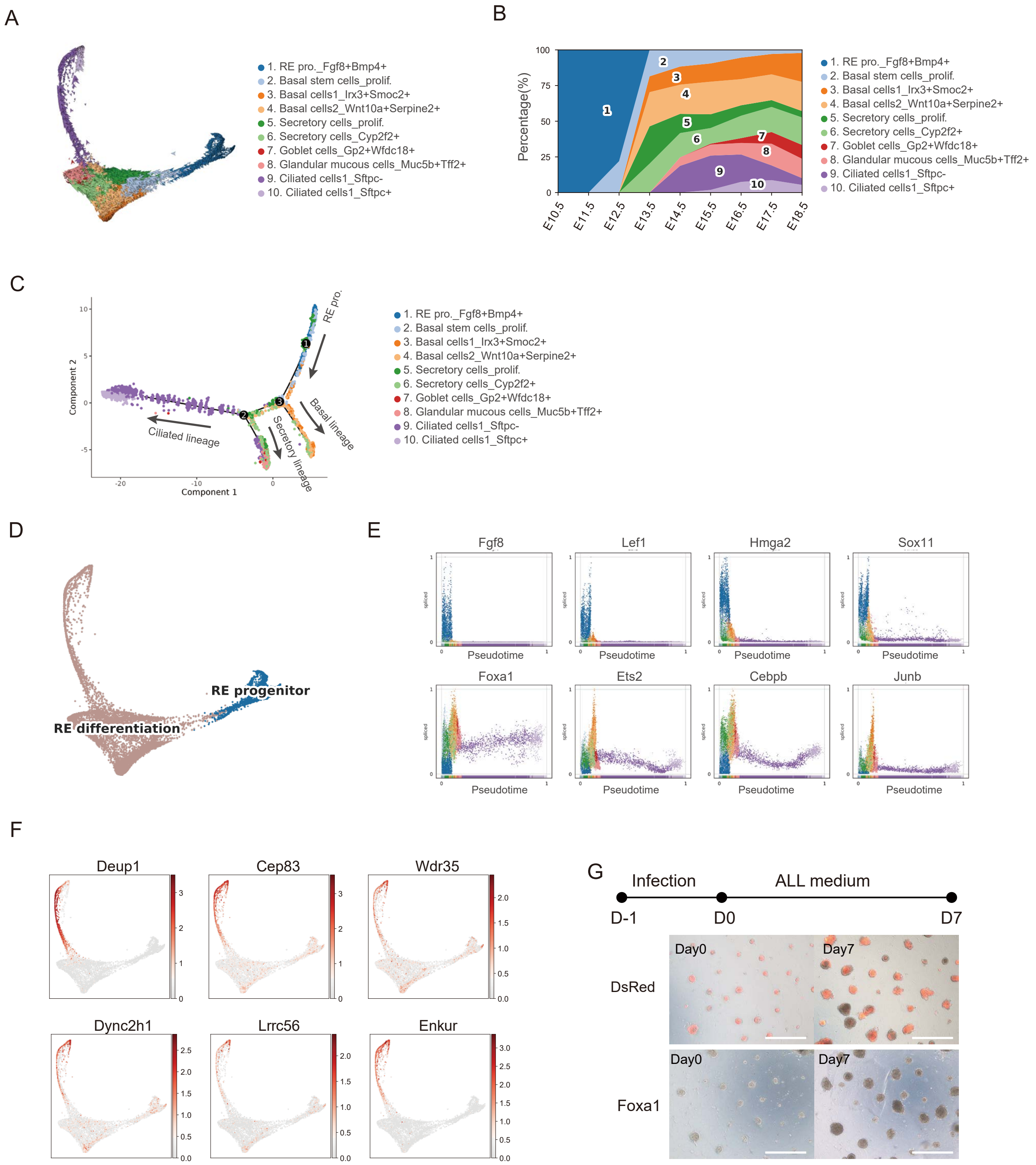

**Fig. S9. Differentiation trajectories and Foxa1-regulated genes in the respiratory epithelium (related to Fig. 5)**

A: RNA velocity of respiratory epithelium clusters is overlaid with velocity streamlines. Velocities were estimated using scVelo's dynamical model based on spliced and unspliced transcript ratios (see Methods).

B: Stacked plot illustrating the fraction of each cell type of RE across embryonic stage.

C: Independent trajectory reconstruction using Monocle2 (DDRTree) showing concordant differentiation paths (details in Methods).

D: PHATE graph illustrating the RE progenitors and downstream RE differentiation states.

E: Scatterplots of normalized spliced transcript abundance of selected genes along pseudotime. *Lef1*, *Hmga2*, *Fgf8* and *Sox11* peak in the RE progenitor state; *Foxa1*, *Ets2*, *Cebpb* and *Junb* are upregulated during the differentiation. Cell colors correspond to panel (C).

F: PHATE projection showing the expression of Foxa1 target genes identified by pySCENIC (corresponding to the red-highlighted genes in Fig. 5F).

G: Representative images of morphologies for the Ctrl (DsRed) and *Foxa1* overexpression cultures. Images are representative of at least three independent experiments. Scale bars, 100 μm.

### Figure S10 related to Fig.6

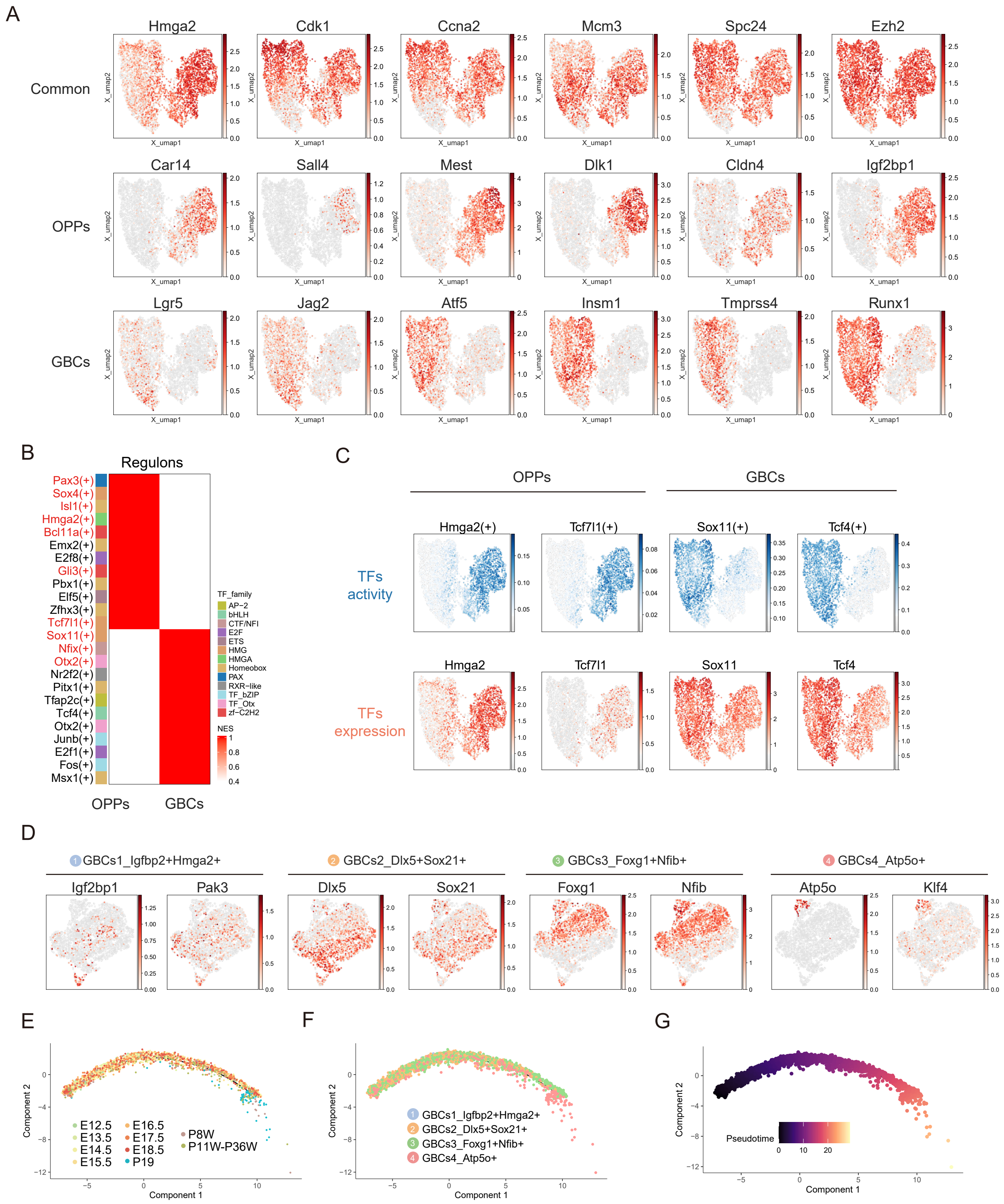

**Fig. S10. Differential characteristics of OPPs and GBCs and the heterogeneity of GBCs (related to Fig. 6)**

A: UMAP visualization of genes shared by OPPs and GBCs (top), OPP-special genes (middle) and GBC-special genes (bottom). Representative genes are selected from DEGs in Fig. 6D.

B: Heatmap of regulon activity (AUC scores) in OPPs versus GBCs by pySCENIC. Regulons are row-z-score normalized. TF families are annotated on the left. Regulons were filtered for measurable activity (see Methods).

C: UMAP showing the TF activity (blue, AUC score) and gene expression (red, log-normalized) for OPPs-specific (*Hmga2* and *Tcf7l1*), and GBCs-specific (*Sox11* and *Tcf4*). SCENIC-generated TF activity, represented by AUC score, reflects the co-expression strength of TF and its target genes.

D: UMAP visualization of differentially expressed genes among four subtypes of GBCs. Genes are selected from DEGs in Fig. 6G.

E: Trajectory reconstruction of GBCs using Monocle2 (DDRTree), colored by embryonic stage to display temporal ordering.

F: The Monocle2 trajectory colored by transcriptional subtypes of GBCs.

G: The Monocle2 trajectory colored by pseudotime. Cells on the left side of the tree (dark colors) correspond to earlier, less differentiated states.
